# Arabidopsis ERF4 and MYB52 Transcription Factors Play Antagonistic Roles in Regulating Homogalacturonan De-methylesterification in Seed Coat Mucilage

**DOI:** 10.1101/2020.01.21.914390

**Authors:** Anming Ding, Xianfeng Tang, Linhe Han, Jianlu Sun, Angyan Ren, Jinhao Sun, Zongchang Xu, Ruibo Hu, Gongke Zhou, Yingzhen Kong

**Author notes:** These authors contributed equally to this work (Anming Ding, Xianfeng Tang). The author responsible for distribution of materials integral to the findings presented in this article in accordance with the policy described in the Instructions for Authors (www.plantcell.org) is Yingzhen Kong,.

## Abstract

The Arabidopsis (*Arabidopsis thaliana*) seed coat mucilage is a specialized cell wall with pectin as its major component. Pectin is synthesized in the Golgi apparatus with homogalacturonan fully methylesterified, but it must undergo de-methylesterification by pectin methylesterase (PME) after being secreted into the cell wall. This reaction is critical for pectin maturation, but the mechanisms of its transcriptional regulation remain largely unknown. Here, we show that the Arabidopsis ERF4 transcription factor positively regulates pectin de-methylesterification during seed development and directly suppresses the expression of *PME INHIBITOR13* (*PMEI13*), *14*, *15* and *SUBTILISIN-LIKE SERINE PROTEASE 1.7* (*SBT1.7*). The *erf4* mutant seeds showed repartitioning of mucilage between soluble and adherent layers as a result of decreased PME activity and increased degree of pectin methylesterification. ERF4 physically associates with and antagonizes MYB52 in activating *PMEI6*, *14* and *SBT1.7* and MYB52 also antagonizes ERF4 activity in the regulation of downstream targets. Gene expression studies revealed that ERF4 and MYB52 have opposite effects on pectin de-methylesterification. Genetic analysis indicated that the *erf4-2 myb52* double mutant seeds show mucilage phenotype similar to wild-type. Taken together, this study demonstrates that ERF4 and MYB52 antagonize each other’s activity to maintain the appropriate degree of pectin methylesterification, expanding our understanding of how pectin de-methylesterification is fine-tuned by the ERF4-MYB52 transcriptional complex in the seed mucilage.

**One-sentence summary:** Arabidopsis ERF4 and MYB52 transcription factors interact and play antagonistic roles in regulating homogalacturonan de-methylesterification related genes in the seed coat mucilage.

## INTRODUCTION

During seed development, some myxospermous species such as Arabidopsis (*Arabidopsis thaliana*) accumulate a large quantity of complex pectinaceous polysaccharides (mucilage) in the apoplast of seed coat epidermal cells or mucilage secretory cells (MSCs). In the Arabidopsis mature seeds, mucilage is compacted between the primary cell wall and the volcano-shaped secondary cell wall called columella (Francoz et al., 2015; Western et al., 2000). When mature dry seeds are imbibed, the rehydrated mucilage ruptures the racial primary cell wall and expands rapidly to form a gelatinous halo that encapsulates the seed. The functional role of the mucilage polysaccharides remains unclear, but this layer has adhesive properties and possibly serves as a water-reservoir, which might facilitate seed dispersion and germination (Francoz et al., 2015; Western et al., 2000).

Mucilage is composed mostly of pectins (Macquet et al., 2007). The Arabidopsis seed coat mucilage provides an excellent model system for studying the biosynthesis, secretion, modification and, critically, the transcriptional regulation of pectins (Arsovski and Behavior, 2010; Francoz et al., 2015; Haughn and Western, 2012). One of the advantages of this model is that pectins can be easily detected using the histochemical stain ruthenium red (RR). The Arabidopsis seed mucilage has been shown to be composed of two layers, termed the water-soluble outer layer and the adherent inner layer (Western et al., 2001). Structural analysis reveals that both layers are composed primarily of pectic polysaccharide rhamnogalacturonan I (RG-I), whereas homogalacturonan (HG) and RG-II are minor components (Golz et al., 2018; Macquet et al., 2007). In the inner layer, cellulose and hemicellulose can also be found, contributing to the tight attachment of the adherent layer to the seed coat, whereas the soluble outer layer is easily lost (Griffiths et al., 2016; Sullivan et al., 2011; Willats et al., 2001a).

Pectins are synthesized in the Golgi apparatus and secreted to the apoplast with as much as 80% of the HG galacturonic acid (GalA) residues being methylesterified, and its de-methylesterification is processed by two kinds of cell wall proteins, termed pectin methylesterase (PME) and pectin methylesterase inhibitor (PMEI) (Pelloux et al., 2007; Wolf et al., 2009). The free carboxylic acid groups resulting from HG de-methylesterification by PMEs can form “egg-box” structures in cross-linking with Ca^2+^, thereby influencing the interactions between HG molecules, and thus affecting cell wall rigidity. In turn, PMEI can impede this process by physically interacting with PME at a 1:1 ratio (Levesque-Tremblay et al., 2015a; Senechal et al., 2015; Wolf et al., 2009). Identification and characterization of Arabidopsis mutants with impaired mucilage extrusion and repartition phenotypes have demonstrated complex mechanisms of pectin structure modifications. For example, establishing the correct degree of methylesterification (DM) is supposedly essential for mucilage extrusion and repartition upon hydration. Function defects of genes such as *PMEI6*, *PMEI14*, *PME58*, *SBT1.7* and *FLY1* are assured to affect the DM, and thereby influencing mucilage structure and organization (Rautengarten et al., 2008; Saez-Aguayo et al., 2013; Turbant et al., 2016; Voiniciuc et al., 2013). In the *pmei6* seeds, methylesterified HG is absent due to high PME activity, leading to mucilage release obstruction (Saez-Aguayo et al., 2013). A similar but less severe phenotype was detected for *pmei14* seeds (Shi et al., 2018). On the contrary, *PME58* promotes HG de-methylesterification and, in the *pme58* seeds, an increase in the DM is attributed to a decrease in PME activity (Turbant et al., 2016). However, no evidence was found for interactions between PME58 and PMEI6 or PMEI14, indicating that more PMEs and PMEIs could participate in mucilage maturation, because both PME and PMEI constitute large multigene families (Turbant et al., 2016; Wang et al., 2013). The mucilage extrusion phenotype was also detected in the *sbt1.7* and *fly1* seeds (Rautengarten et al., 2008; Voiniciuc et al., 2013). However, PMEI6 and SBT1.7 may target different PMEs, given that the *pmei6 sbt1.7* double mutant presents additive phenotypes (Saez-Aguayo et al., 2013). The E3 ubiquitin ligase FLY1 is supposed to regulate pectin de-methylesterification by recycling PMEs in the MSC endomembrane system (Voiniciuc et al., 2013).

Although enzymes involved in the regulation of pectin de-methylesterification (either positively or negatively) have been reported, the mechanisms of their transcriptional regulation are largely unknown. At present, only three transcriptional regulators, LUH/MUM1, STK and MYB52 of pectin methylesterification modification were identified in the seed coat mucilage. LUH/MUM1 promotes the expression of *PMEI6* and *SBT1.7* (Huang et al., 2011; Walker et al., 2011). However, LUH/MUM1 also seems to be a positive regulator of pectin de-methylesterification as PME activity is reduced in *luh/mum1* seeds (Saez-Aguayo et al., 2013). STK was found negatively regulating pectin de-methylesterification through direct activation of *PMEI6*. Therefore, the *stk* seeds present similar phenotypes to *pmei6*. STK also represses the expression of *SBT1.7* (Ezquer et al., 2016). What’s more, LUH/MUM1 and STK can antagonize each other’s function since each represses the other’s activity. Our recent work reported that MYB52 transcription factor negatively regulates pectin de-methylesterification in the seed mucilage by directly activating *PMEI6*, *PMEI14* and *SBT1.7*. Thus in the *myb52* seeds, an increase in PME activity and a decrease in DM were observed, leading to a condensed “thin” inner layer (Shi et al., 2018). These results suggest that mucilage methylesterification modifications underlie the regulation of a complex regulatory network.

The Arabidopsis AP2/ERF superfamily was divided into AP2, ERF, RAV families and a soloist gene (Nakano et al., 2006). To date, the only identified member of the AP2 family is *AP2*, which likely plays an indirect role in mucilage production, considering that the MSCs cannot be differentiated in the *ap2* seeds (Jofuku et al., 1994; Western et al., 2001). *<u>E</u>THYLENE <u>R</u>ESPONSIVE <u>F</u>ACTOR4* (*ERF4*) belongs to the ERF family, Group VIII (B-1a), and all eight genes, *ERF3*, *ERF4* and *ERF7*~*12* from this subgroup were proved to be transcriptional repressors (Fujimoto et al., 2000; Koyama et al., 2013; Maruyama et al., 2013; Yang et al., 2005). *ERF4* has been shown to have diverse functions in plant growth and development, such as modulating ethylene, abscisic acid (ABA), jasmonic acid (JA) and gibberellic acid (GA) responses (McGrath et al., 2005; Yang et al., 2005; Zhou et al., 2016), operating in the progression of leaf senescence (Koyama et al., 2013; Riester et al., 2019), enhancing resistance to pathogens (McGrath et al., 2005), promoting internode elongation (Zhou et al., 2016), and also playing important roles in nutrition stress (Liu et al., 2017). *ERF4* is expressed ubiquitously in Arabidopsis (Winter et al., 2007), suggesting that it has other functions in addition to those mentioned above. For example, its regulatory role in cell wall biology has not been reported yet.

Here, we provide evidence that *ERF4* positively regulates pectin de-methylesterification in the seed mucilage. We find a repartitioning of seed mucilage polysaccharides in the *erf4* seeds under vigorous shaking conditions which can be attributed to an increase in DM. ERF4 binds to the *cis*-regulatory elements and directly suppresses the expression of *PMEI13*, *14*, *15* and *SBT1.7*. Moreover, ERF4 and MYB52 were found to interact, and play completely opposite roles in pectin de-methylesterification by antagonizing each other’s transcriptional activity, which is confirmed by gene expression and genetic evidence. Overall, these findings provide a fine-tuned mechanism where ERF4 and MYB52 antagonistically control mucilage modification related genes.

## RESULTS

### Expression pattern of *ERF4* is correlated with seed mucilage deposition

Previous studies revealed that *ERF4* is expressed in various tissues with its transcripts accumulating predominantly in developing seeds (Winter et al., 2007) (Supplemental Figure 1A). Our re-analysis of the microarray datasets for Laser Capture Microdissected seed samples (GSE12404) identified a significant up-regulation of *ERF4* during seed coat development stages (Hu et al., 2016). To confirm these findings, we investigated the expression pattern of *ERF4* by qRT-PCR in developing seeds at 4, 7, 9, 11 and 13 days-post-anthesis, (DPA) as well as in major Arabidopsis organs. We observed that *ERF4* transcripts were detected in all examined tissues. In particular, *ERF4* showed a strong expression in developing siliques with the highest level found in 13 DPA seeds, which is equivalent to the mature green embryo stage (Figure 1A). This expression pattern coincides with the period for mucilage polysaccharides production in the seed coat (Francoz et al., 2015).

**Figure 1.**
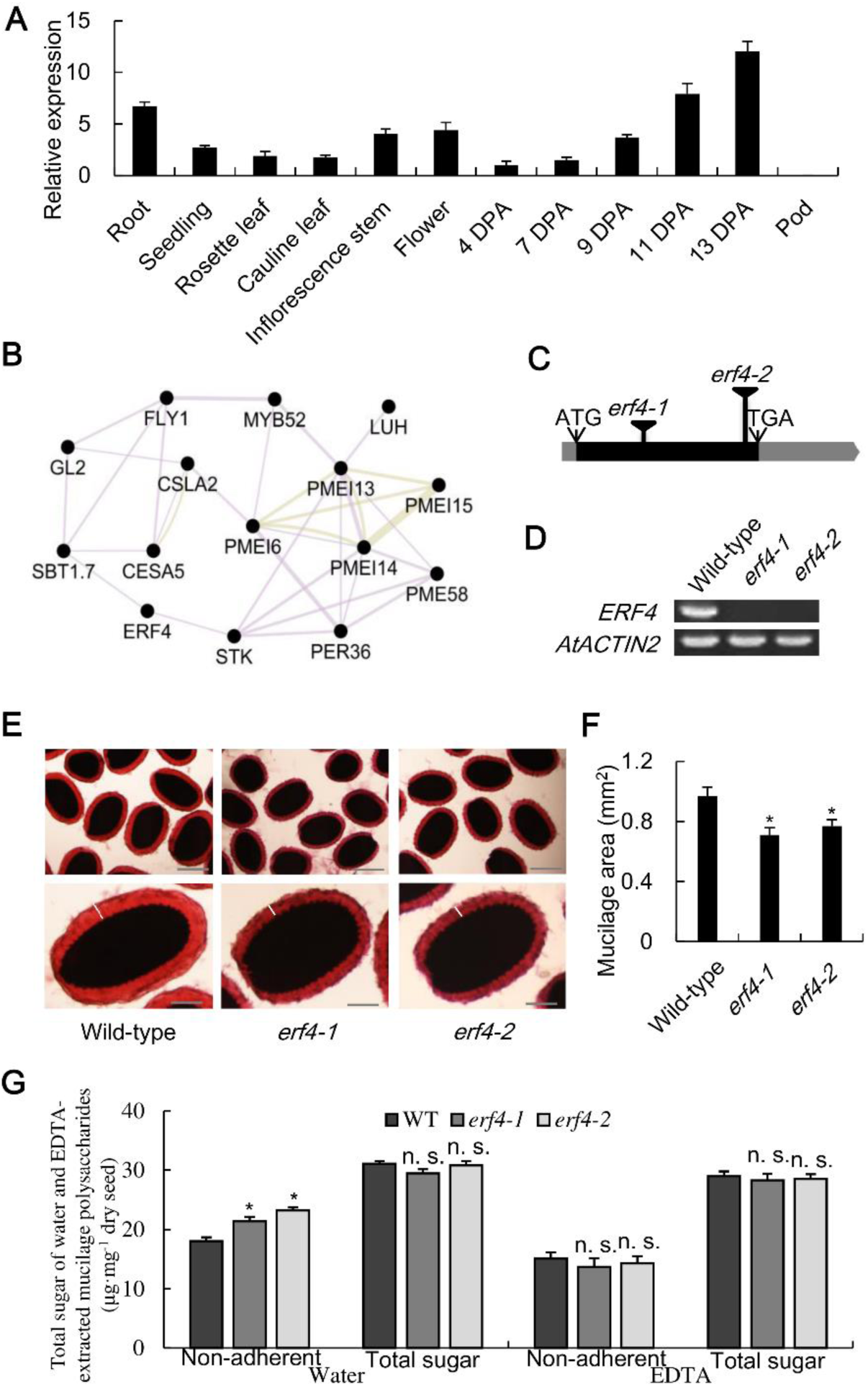
Expression analysis of *ERF4* and the *erf4* mutants present a mucilage phenotype under vigorous shaking conditions. **(A)** Relative expression of *ERF4* in developing siliques at 4, 7, 9, 11 and 13 DPA and major Arabidopsis organs with qRT-PCR analysis. Gene expressions were measured using the reference gene *ACTIN2*. Values correspond to means ± SD of three replicates for each sample. The expression level at 4 DPA was set as 1. **(B)** Co-expression network of *ERF4* with known mucilage genes based on GeneMANIA. *GL2*, Glabra2; *LUH/MUM1*, Mucilage-modified 1; *MYB52*, MYB domain protein 52; *STK*, SEEDSTICK; *FLY1*, Flying Saucer 1; *CSLA2*, Cellulose synthase-like A2; *CESA5*, Cellulose synthase 5; *SBT1.7*, Subtilisin-like serine protease 1.7; *MUCI10*, Mucilage-related 10; *PER36*, Peroxidase 36; *PMEI6, 13, 14, 15*, Pectin methylesterase inhibitor 6, 13, 14, 15; *PME58*, Pectin methylesterase 58. Co-expression was indicated with purple lines, while proteins share similar domains with yellow green. **(C)** Gene model of *ERF4* with T-DNA insertion sites. The black line shows coding region (CDS), while gray lines represent non-coding upstream and downstream regions. **(D)** Semi-quantitative RT-PCR was performed on cDNA from siliques of wild-type and *erf4* mutants with primers flanking the full-length CDS. *ACTIN2* was amplified as the loading control. **(E)** RR staining of adherent mucilage (AM) for wild-type and *erf4* seeds. Mature dry seeds were shaking in distilled water with 0.01% RR at 200 rpm for 1 h before being photographed with an optical microscope. Bars = 500 μm and 100 μm, respectively. **(F)** Area of AM layers. Values are average area ± SD of at least 20 seeds which were measured with the optical microscope from the top to the bottom of the red halo (white lines in **(E)**). **(G)** Comparison of total sugar contents in AM layers and whole mucilage of wild-type and *erf4* seeds extracted by shaking at 200 rpm for 1 h in water or EDTA. *, *P* < 0.05; n. s., not significant, Student’s t-test.

To obtain more detailed information on its spatial and developmental expression profile, a *proERF4::GUS* construct was generated and transformed in wild-type plants. *GUS* expression was detected in vascular tissues, guard cells and main roots of 5-day old seedlings (Supplemental Figure 2A to C). In mature plants, GUS activity was detected in root tips, inflorescence internodes, mature rosette and cauline leaves (Supplemental Figure 2D to G). In blooming flowers, GUS staining was observed in petals, stigma and pollen (Supplemental Figure 2H). The GUS signal was also notably detected in 4 DPA siliques and MSCs of developing seeds (Supplemental Figure 2I to K). Taken together, these results confirm the expression of *ERF4* in testas of developing seeds when mucilage polysaccharides are produced during 7~13 DPA of embryogenesis. In addition, using Cytoscape v3.7.1 software equipped with the GeneMANIA tool (Warde-Farley et al., 2010), *ERF4* was predicted to be co-expressed with multiple mucilage genes (Figure 1B). These results suggest that *ERF4* might be involved in the regulation of mucilage deposition in the seed coat.

### Area of mucilage capsule is reduced in the *erf4* mutants

Two independent T-DNA insertion lines, *erf4-1* and *erf4-2* (*erf4* hereafter, unless specified) were isolated in the background of Col-0 ecotype (Figure 1C). The transcripts of *ERF4* in *erf4* homozygous were detected by RT-PCR. Results showed that both lines are null mutants for *ERF4* (Figure 1D).

To determine whether *ERF4* functions in seed mucilage production, mature dry seeds were examined by RR staining. Upon imbibition in distilled water with moderate shaking, no significant difference was observed (Supplemental Figure 3A and B). However, when shaken at a frequency of 200 rpm for 1 h, the *erf4* seeds showed thinner mucilage halos and the area of adherent mucilage was reduced by ~30% compared to wild-type (Figure 1E and F). In addition, when *erf4-2* was transformed with functional *ERF4*, driven by the cauliflower mosaic virus (CaMV) *35S* promoter (*35S::ERF4*), the reduced adherent mucilage phenotype was complemented (Supplemental Figure 4A and B). These results support that the deletion of *ERF4* is responsible for the mucilage defect.

We also examined seed coat morphology to clarify effects of the *ERF4* mutation on seed coat development. Scanning electron microscopy of the mature seeds’ surface showed that the volcano-shaped collumella and the radial wall were similar between wild-type and *erf4-2* seeds (Supplemental Figure 3C and D). Developing seeds (at 7 and 10 DPA) were sectioned and stained to determine if the seed mucilage deposition was affected. The results showed that the shape of the MSCs was similar between *erf4-2* and wild-type, and the deposition of mucilage was also unaffected (Supplemental Figure 3E). Altogether, these results revealed that the *erf4-2* seed coat morphology is unaltered compared with wild-type, but its adherent mucilage layer seems to be loose and can be extracted more easily than wild-type in water, indicating that the *ERF4* mutation might affect pectin structure and organization during mucilage maturation.

### Mucilage partitioning is modified in *erf4* seeds

We have observed that the area of *erf4* adherent layer was smaller than that of wild-type seeds. To testify whether the distribution of mucilage layers was affected, water-soluble mucilage (SM) and adherent mucilage (AM) were sequentially extracted from *erf4* and wild-type mature seeds in deionized water (Supplemental Figure 3F and G), and analyzed for their sugar content and composition (Supplemental Table 1). As the major component of Arabidopsis seed mucilage is RG I, compositional analysis showed that both SM and AM layers of wild-type and *erf4* consisted predominantly of rhamnose (Rha) and GalA at a molar ratio close to 1. A small quantity of other monosaccharides was also detected. However, in the *erf4* SM layer, the content of Rha and GalA was increased by ~20% compared to that of wild-type seeds. In contrast, in the AM layer, the content of Rha and GalA decreased by 21 and 50% (in *erf4-1* and *erf4-2*, respectively), compared to that in wild-type.

As expected, the amount of mucilage in the *erf4* AM layer was reduced to ~60% of that in wild-type, whereas in its SM layer the quantity of mucilage sugars was increased by ~20% (Figure 1G). The *erf4* seeds thus showed a re-distribution of the SM and AM layers. However, the total amount of mucilage sugars across the two fractions was nearly unaltered between *erf4* and wild-type (Figure 1G). Moreover, no appreciable differences were noted in the total content of Rha (~43, ~39 and ~40% in *erf4-1*, *erf4-2* and wild-type seed mucilage, respectively) or GalA (~48, ~51 and ~52% in *erf4-1*, *erf4-2* and wild-type seed mucilage, respectively). The above results suggest that the mutation in *ERF4* does not affect the biosynthesis of seed coat mucilage but affects the modifications of mucilage structure.

### *ERF4* mutation affects HG methylesterification in adherent mucilage

The DM is believed to have impacts on mucilage physical properties, and thus can affect mucilage extrusion and repartition. To establish whether DM is altered in the *erf4* seed mucilage, we determined DM indirectly by quantification of the formaldehyde produced from methanol by alcohol oxidase (Klavons and Bennett, 1986). In *erf4-1* and *erf4-2*, DM was increased by ~10% and ~19% compared to that in wild-type, respectively (Figure 2A). Since the methanol released from seed mucilage is supposed to be derived from methylesterified HGs (Ezquer et al., 2016), our results suggest that *ERF4* might play a role in promoting HG de-methylesterification in the seed mucilage.

**Figure 2.**
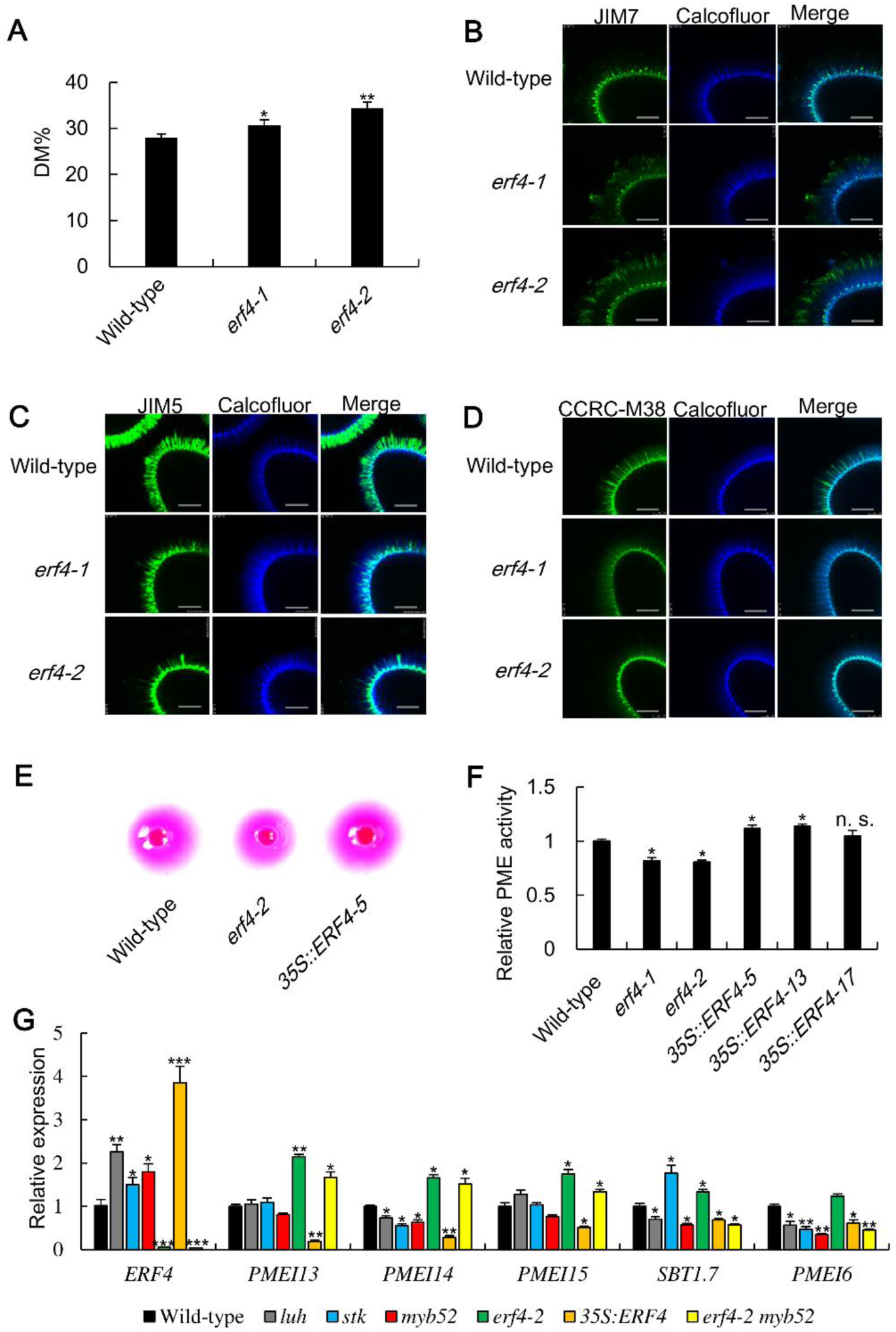
ERF4 positively regulates HG de-methylesterification by promoting PME activity. **(A)** Degree of pectin methylesterification (DM%) of wild-type and *erf4* seed mucilage. Error bars represent SD values of three biological experiments. **(B)** to **(D)** Immunodetection experiments were performed on adherent mucilage released from whole seeds. Optical sections of AM were visualized by Confocal microscopy. HGs of different methylesterified state were recognized with JIM7 **(B)**, JIM5 **(C)** and CCRC-M38 **(D)** antibodies, respectively. Calcofluor white was used to label cellulose which were shown as blue rays in the sections. Bars = 50 μm. **(E)** PME activities of total protein extracts from whole seeds of wild-type, *erf4-2* and *35S::ERF4* were visualized in gel diffusion analysis. **(F)** Relative PME activity was measured according to gel diffusion and was normalized to average wild-type activity (=1). Error bars represent SD values of three biological experiments. **(G)** Expression pattern of *ERF4*, *PMEI13*, *PMEI14*, *PMEI15*, *PMEI6* and *SBT1.7* in mutants, icluding *luh*, *stk*, *myb52*, *erf4-2*, *35S::ERF4* and *erf4-2 myb52*, compared to that in wild-type. Gene expressions were measured using the reference gene *ACTIN2*. The values represent means of three biological replicates ± SD. The expression level for each gene in wild-type seeds was set as 1. *, *P* < 0.05; **, *P* < 0.01; ***, *P* < 0.001, Student’s t-test.

The epitopes presented in the seed coat surface can be recognized and labeled by certain antibodies through immunolabeling experiments. For example, JIM5, JIM7 and CCRC-M38 antibodies can recognize poorly methylesterified HGs, highly methylesterified HGs and non-esterified HGs, respectively (Macquet et al., 2007; Willats et al., 2001b). To further determine whether *ERF4* affects DM, whole-seed immunolabeling was performed on wild-type and *erf4* seeds. Upon rehydration of mature dry seeds soluble mucilage was easily lost, resulting in immunolabeling of mainly adherent mucilage. In wild-type seeds, the JIM7 signal was located throughout the AM, and a significant increase in labelling signal was observed for both *erf4-1* and *erf4-2* (Figure 2B). In turn, JIM5 labelling showed a notable decrease of less methylesterified HGs along the ray structures in *erf4* compared to wild-type (Figure 2C). When immunolabeling was performed with CCRC-M38, the labelling was also significantly reduced (Figure 2D). De-methylesterification is supposed to facilitate the formation of calcium-mediated cross-links between neighboring HG molecules, which can be recognized by 2F4 antibody (Willats et al., 2001a). Our immunolabeling analysis showed that the labelling intensity was strongly reduced in *erf4-2* seeds compared to wild-type, suggesting a decreased amount of this cross-linking in mutants (Supplemental Figure 5). Collectively, the mutation in *ERF4* inhibits pectin de-methylesterification in the seed mucilage.

### PME activity is inhibited in the *erf4* seeds

To identify whether the increased DM in *erf4* seeds was a result of decreased PME activity, total proteins were extracted from whole seeds of wild-type, *erf4*, *erf4-2*:*35S::ERF4* and *35S::ERF4* overexpressors, and PME activity was determined as previously described by Lekawska-Andrinopoulou et al. (2013) with the average wild-type PME activity being normalized to 100% (=1). The results showed that PME activity was reduced by ~13% and ~17% in the *erf4-1* and *erf4-2* seeds, whereas it was elevated by 5~14% in *35S::ERF4* lines compared to wild-type (Figure 2E and F). As expected, PME activity in *erf4-2*:*35S::ERF4* was not significantly altered (Supplemental Figure 4C). Collectively, ERF4 positively regulates mucilage de-methylesterification by promoting PME activity.

We also observed that, compared to wild-type, the seed size of *35S::ERF4* was increased on average by ~12% in length and ~9% in width, resulting in a significant increase of seed area (Supplemental Figure 6A to C). Furthermore, a significant increase of MSC surface area in *35S::ERF4* seeds was also detected (Supplemental Figure 6D and E). So, even if structure of epidermal cells of the *erf4-2* seeds appears unchanged, overexpression of *ERF4* seems to promote seed coat epidermal cell and seed size.

### ERF4 affects the expression of genes involved in DM modification

PMEI can inhibit PME activity through direct protein-protein interaction, thus we speculate that as a transcription repressor ERF4 might promote PME activity by suppressing certain *PMEI* genes. To test this hypothesis, expression fold change of the *PMEI* gene family (Wang et al., 2013) was investigated by qRT-PCR using RNAs from *erf4-2*, *35S::ERF4* and wild-type developing seeds at 13 DPA. *PMEI* genes including *PMEI6*, *PMEI13*, *PMEI14*, *EDA24*, *AT1G11593*, *AT1G70720*, *AT3G05741*, *AT4G03945* and *AT4G12390* had similar expression pattern, being significantly up-regulated in *erf4-2* and down-regulated in *35S::ERF4* (Figure 2G; Supplemental Figure 7A) compared to that in wild-type seeds. The ERF transcription factors are well known for binding to *cis*-acting elements of GCC-box (GCCGCC) or DRE-motif (CCGAC). Of the 9 *PMEIs*, *PMEI13*, *PMEI14* and *AT3G05741* (termed *PMEI15* hereafter) have predicted GCC-box motifs in their promoter or intron regions (Figure 3A to C). Although *PMEI6* was included, no ERF binding site was found in its promoter, suggesting that ERF4 might negatively regulate its expression indirectly.

**Figure 3.**
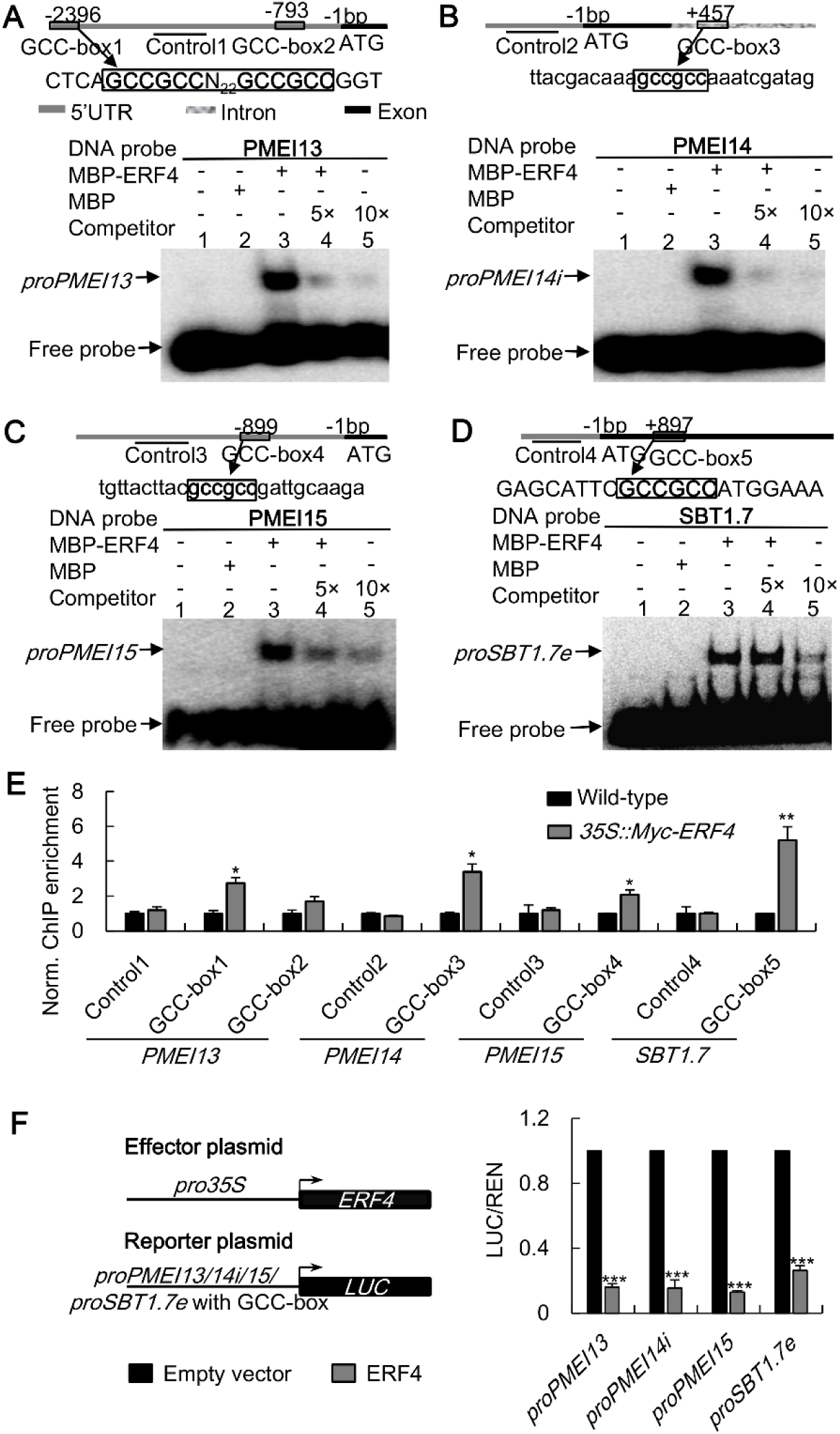
Identification of *PMEI13*, *PMEI14*, *PMEI15* and *SBT1.7* as downstream targets of ERF4. **(A)** to **(D)** EMSA analysis showing that ERF4 binds to the GCC-box motif in the promoters, intron or exon of *PMEI13* **(A)**, *PMEI14* **(B)**, *PMEI15* **(C)** and *SBT1*.*7* **(D)**. Relative nucleotide positions of putative ERF binding site are indicated (with the first base preceding the ATG start codon being assessed as -1). The DNA probes were 5’biotin-labeled fragments containing the GCC-box motif. Competitors were the same fragments but were non-labeled (5 and 10 fold that of the hot DNA probe). MBP-tagged ERF4 fusion protein (MBP-ERF4) was purified, and the MBP tag was used as negative control. The sequences show the GCC-box motif which is highlighted in bold. **(E)** ChIP-qPCR analysis for chromatin extracts from transgenic plants expressing Myc-ERF4 fusion protein and wild-type. Nuclei from developing seeds of Myc-ERF4 transgenic lines and wild-type were immunoprecipitated by anti-Myc antibody. The precipitated chromatin fragments were analyzed by qPCR with primer amplifying these GCC-box containing regions (GCC-box1, -box2, -box3, -box4 and -box5) as indicated in **(A)**, **(B)**, **(C)** and **(D)**. Error bars represent means ± SD from three independent experiments. **(F)** Relative LUC activity analysis showing that ERF4 suppresses the expression of *PMEI13*, *PMEI14*, *PMEI15* and *SBT1.7*. Dual-LUC transient transcriptional activity assays were performed in wild-type Arabidopsis protoplasts. Expression of *ERF4* and the *REN* internal control was driven by the CaMV 35S promoter and that of the LUC reporter gene was driven by promoters of *PMEI13* or *PMEI15*, first intron of *PMEI14* (*PMEI14i*) or exon of *SBT1.7* (*SBT1.7e*) containing GCC-box motif (*proPMEI13*, *proPMEI14i*, *proPMEI15* or *proSBT1.7e::LUC*). LUC/REN represents the relative activity of promoters. Three independent experiments were performed. The values present the means ± SD. *, *P* < 0.05; **, *P* < 0.01; ***, *P* < 0.001, Student’s t-test.

We also examined relative expression levels for genes reported to be associated with PME activity, such as *LUH/MUM1*, *STK*, *MYB52*, *FLY1*, *SBT1.7, PME58* and *UUAT1*. The qRT-PCR results showed that except *SBT1.7* the others had no significant change in expression (Figure 2G; Supplemental Figure 7B). *Cis*-element analysis showed that *SBT1.7* also has one GCC-box motif in its exon region (Figure 3D).

### ERF4 binds the GCC-box motifs in *PMEI13*, *14*, *15* and *SBT1.7*

To test whether ERF4 could bind the GCC-box motifs in *PMEI13*, *14*, *15* and *SBT1.7*, we designed 5’-Biotin labelled probes (Supplementary Table 2) containing the GCC-box motif in electrophoretic mobility shift assays (EMSA) to assess the binding ability of ERF4 to these fragments *in vitro*. As hypothesized, purified MBP-ERF4 fusion protein bound to the GCC-box containing fragments of both *PMEI13*, *14*, *15* and *SBT1.7* (Figure 3A to D, lane 3). When MBP was added alone, no mobility shift was observed (Figure 3A to D, lane 2). Moreover, the binding of ERF4 to these labelled probes was weakened by the addition of an unlabeled competitor (Figure 3A to D, lane 4 and 5).

To assess whether ERF4 binds to these GCC-box motifs *in vivo*, we conducted chromatin immunoprecipitation (ChIP)-qPCR assays with chromatin extracts from developing seeds of wild-type plants and plants overexpressing Myc-tagged ERF4 (*35S::Myc-ERF4*). The presence of ERF4 substantially enhanced detection of the GCC-box containing sequences in the promoters of *PMEI13*, *15*, intron of *PMEI14* (*14i*) and exon of *SBT1.7* (*SBT1.7e*), indicating that ERF4 bound to these sequences *in planta* (Figure 3E). Finally, the transcriptional regulation of ERF4 to these targets was evaluated through the detection of relative LUC activity in wild-type Arabidopsis protoplasts with REN being used as internal control. Effector plasmid *35S::ERF4* and reporter plasmids *proPMEI13*, *14i*, *15* or *SBT1.7e::LUC* were generated. When *35S::ERF4* was co-transformed with either reporter plasmid, the LUC activity was significantly inhibited (Figure 3F). To test the authenticity of ERF4 suppressing *SBT1.7* expression by binding to the GCC-box motif in its exon region, co-transformation of *35S::ERF4* and *promuSBT1.7e* with a mutational GCC-box abolished this binding (Supplemental Figure 8). Together, our data indicate that ERF4 could negatively regulate the expression of *PMEI13*, *14*, *15* and *SBT1.7* by directly binding to the regulatory elements that locate in the promoter, intron or exon region.

### Functions of *PMEI13* and *PMEI15* in seed mucilage maturation

Both *PMEI13*, *14*, *15* and *SBT1.7* are predominantly expressed in developing seeds at either 4 or 7 DPA, as verified in our qRT-PCR analysis (Supplemental Figure 1B to F). The roles of *PMEI14* and *SBT1.7* in seed mucilage maturation have been well established (Rautengarten et al., 2008; Shi et al., 2018). To determine if *PMEI13* and *PMEI15* have similar functions in seed mucilage maturation, seeds of *pmei13* and *pmei15* null mutants were stained with RR. Under vigorous shaking conditions, *pmei13-1* and *pmei13-2* seeds presented thinner AM layers compared with wild-type, whereas the area of mucilage halos of *pmei15-1* and *pmei15-2* seeds showed no significant difference from wild-type (Supplemental Figure 9A to D). Compared to wild-type, DM of seed mucilage decreased significantly in *pmei13-1* seeds, but not in *pmei15-1* seeds (Supplemental Figure 9E). We also performed PME activity assays on *pmei13-1* and *pmei15-1* whole seeds (Supplemental Figure 9F). Results showed that, consistent with the decreased DM, *pmei13-1* presents a significant increase in PME activity compared to wild-type. However, no significant difference was observed between *pmei15-1* and wild-type, suggesting that the function of *PMEI15* might be minimal, if any, in seed mucilage maturation.

### Role of ERF4 in DM modification is mediated by *PMEI13*, *14*, *15* and *SBT1.7*

Genetic interactions between *ERF4* and *PMEI13*, *14*, *15* or *SBT1.7* in modulating DM were studied by generating *erf4-2 pmei13-1*, *erf4-2 pmei14*, *erf4-2 pmei15-1* and *erf4-2 sbt1.7* double mutant lines. In line with our findings that *PMEI13*, *14*, *15* are downstream targets of ERF4 with enhanced expression in the *erf4-2* seeds, both *pmei13-1* and *pmei14* single mutations, but not the *pmei15-1* single mutation, partially rescued the mucilage phenotype and the reduction of PME activity in *erf4-2* seeds (Supplemental Figure 9C to F). However, *erf4-2 sbt1.7* seeds resembled *sbt1.7* single mutant in mucilage extrusion phenotype and PME activity. Consistent with the observed phenotype, the DM of *erf4-2 pmei13-1* and *erf4-2 pmei14* double mutant seeds was also partially restored to wild-type levels compared to *erf4-2*, whereas *erf4-2 sbt1.7* seeds had a compromised DM which was between *erf4-2* and *sbt1.7*, but was still significantly decreased compared to wild-type (Supplemental Figure 9E).

### ERF4 physically interacts with MYB52

Our previous study has shown that MYB52 could directly activate *PMEI6*, *14* and *SBT1.7* (Shi et al., 2018). In this work, we have demonstrated that ERF4 directly suppresses *PMEI14* and *SBT1.7*. These findings led us to explore the relationships between the ERF4 and MYB52 transcription factors. Firstly, qRT-PCR was performed to determine the change of expression levels for *ERF4* and *MYB52* in *myb52* and *erf4-2* seeds, respectively. Whereas expression of *MYB52* in *erf4-2* was unaltered (Supplemental Figure 7B), *ERF4* expression was increased in *myb52* (Figure 2G). We next examined *cis*-acting elements in the *ERF4* promoter and found three MYB binding sites (MBS). However, MYB52 did not directly bind to these MBS motifs in our EMSA and ChIP assays (data not shown). These results suggest that ERF4 and MYB52 do not directly regulate each other at a transcriptional level.

Nevertheless, ERF4 and MYB52 might still regulate each other’s function through direct protein-protein interaction. To verify this hypothesis, we performed a GST pull-down assay to verify whether these two transcription factors can interact. As expected, purified MBP-ERF4 fusion protein was pulled down when incubated with GST-tagged MYB52 (GST-MYB52) using anti-MBP antibodies (Figure 4A). No band was detected in the immunoblotting analysis when pull-down was executed with the GST tag alone, suggesting that the interaction between ERF4 and MYB52 is specific. To determine whether ERF4 and MYB52 interact *in vivo*, co-immunoprecipitation (co-IP) assays were performed using Arabidopsis protoplasts transformed with *35S::MYB52-Myc* and *35S::ERF4-Flag* constructs. The results showed that MYB52-Myc was identified using the anti-Myc antibody when the protein extracts were immunoprecipitated with anti-flag antibody, indicating that ERF4 and MYB52 interact in plant cells (Figure 4B). Lastly, to further assure the interaction between ERF4 and MYB52 *in planta*, we conducted a bimolecular fluorescence complementation (BiFC) assay in Arabidopsis protoplasts (Figure 4C). ERF4-YC and YN-MYB52 co-expression could reconstitute YFP fluorescence. In contrast, co-expression of ERF4-YC with the YFP N-terminus or YFP C-terminus with YN-MYB52 or the YFP C-terminus and N-terminus alone did not show any fluorescence signal. Moreover, the BiFC results also showed that the interaction between ERF4 and MYB52 occurs in the nucleus.

**Figure 4.**
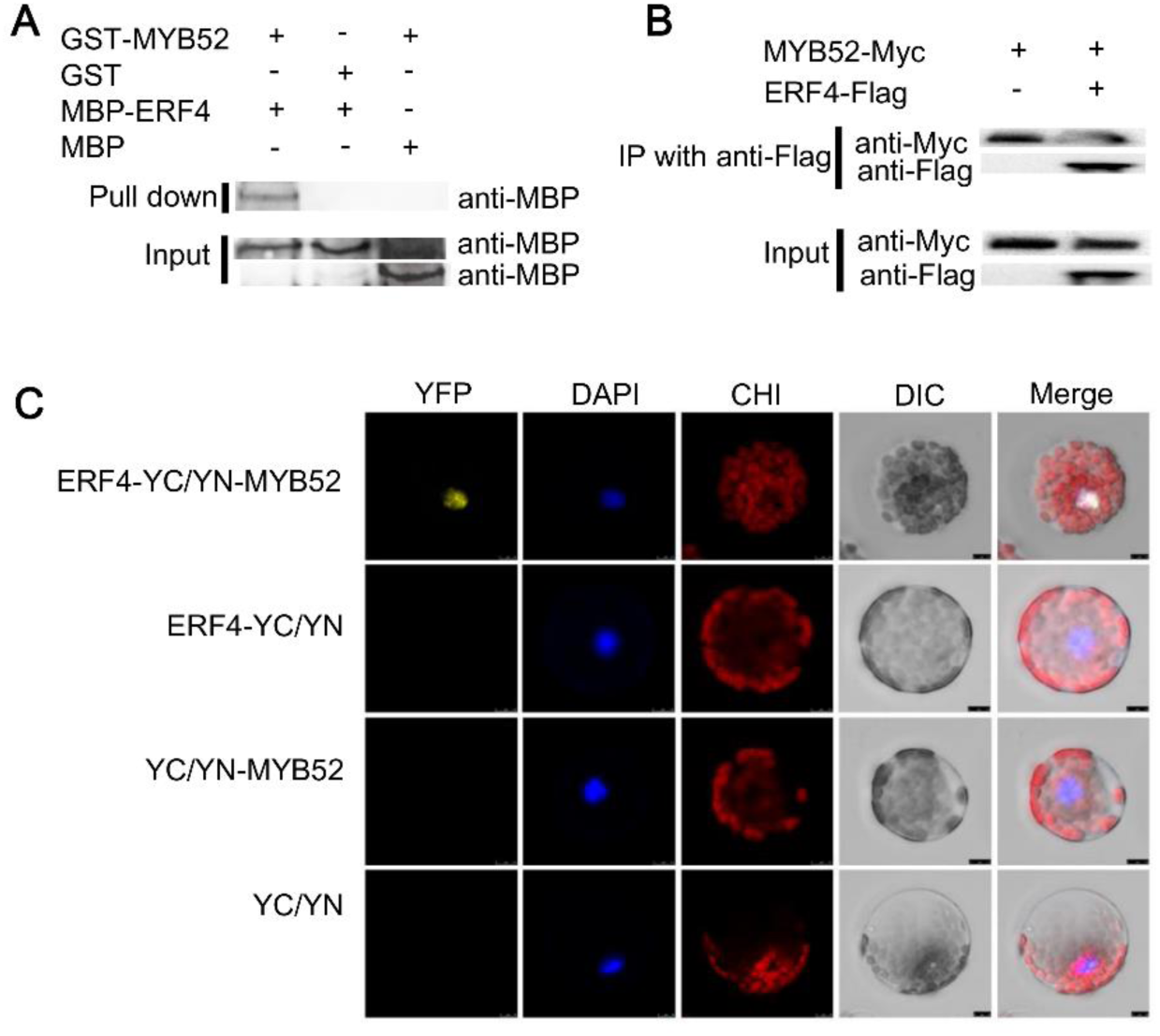
ERF4 physically interacts with MYB52. **(A)** Pull-down assay of ERF4 interaction with MYB52. GST tagged MYB52 fusion protein (GST-MYB52) or GST alone were incubated with MBP tagged ERF4 fusion protein (MBP-ERF4) in GST beads. MBP-ERF4 but not MBP was pulled down by the beads containing GST-MYB52. MBP tag alone incubated with GST-MYB52 in GST beads was used as negative control. **(B)** In vivo co-immunoprecipitation assay of ERF4 and MYB52 interaction. Flag tagged ERF4 (ERF4-Flag) and Myc tagged MYB52 (MYB52-Myc) were overexpressed in Arabidopsis protoplasts. Flag antibody was used for immunoprecipitation analysis, and Myc antibody was used for immunoblot analysis. The band detected by Myc antibody in the precipitated protein sample indicates the physical interaction between ERF4 and MYB52. **(C)** Interaction of ERF4 and MYB52 in Arabidopsis protoplast. Representative cells were shown which were imaged by confocal laser scanning microscopy. The yellow fluorescence for YFP was only detected in Arabidopsis protoplasts of ERF4-YC and YN-MYB52 interaction. DAPI, nucleus labeled by DAPI in blue fluorescence; CHI, chloroplast auto-fluorescence in red signal; DIC, bright field. Bars = 10 μm.

### ERF4 and MYB52 antagonize each other’s activity in regulating pectin de-methylesterification related genes

To test whether the ERF4-MYB52 interaction affects the binding of ERF4 to *PMEI13*, *14*, *15* and *SBT1.7*, EMSA assays were performed with purified MBP-ERF4 and GST-MYB52 fusion proteins and biotin labelled probes that only contain the GCC-box motif. Results showed that ERF4 alone bound to the DNA probes containing GCC-box (Figure 5A to D, lane 3), while MYB52 did not (Figure 5A to D, lane 2), and when the amount of MYB52 was added increasingly, the binding of ERF4 to these probes gradually weakened (Figure 5A to D, lane 4 to 6). To investigate how the ERF4-MYB52 interaction affects their transcriptional activity, an effector plasmid of *35S::MYB52* was additionally constructed, and dual-LUC transient transcriptional activity assays were performed in Arabidopsis protoplasts (Figure 5E). *35S::MYB52* co-transformed with either reporter plasmid showed no influence on LUC activity, indicating that MYB52 did not activate promoters containing the GCC-box motif. When *35S::ERF4* and *35S::MYB52* were co-transformed together with either reporter plasmid, the LUC activity was significantly recovered in comparison with co-transformations without *35S::MYB52*, indicating that the transcriptional suppression of ERF4 to its downstream genes was inhibited by MYB52.

**Figure 5.**
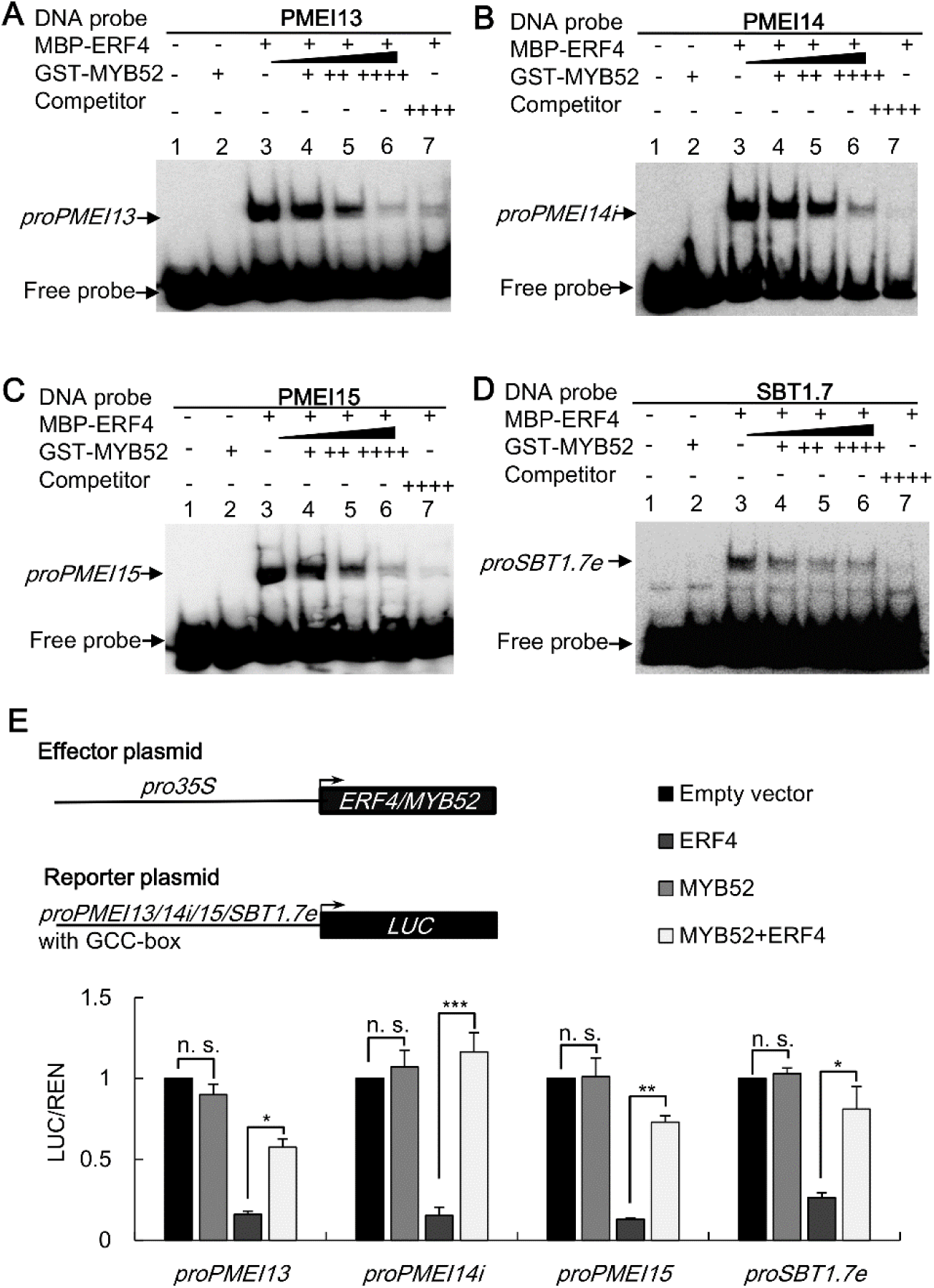
MYB52 interfered ERF4 function in regulating its downstream targets. **(A)** to **(D)** EMSA results showing that MYB52 did not bind to probes containing GCC-box from *PMEI13* **(A)**, *PMEI14* **(B)**, *PMEI15* **(C)** and *SBT1.7* **(D)** (lane 2), but ERF4 did bind to those motifs (lane 3). MYB52 interfered the binding of ERF4 to these promoters (lane 4 to 6). The DNA probes were 5’biotin-labeled fragments of the gene promoters containing the GCC-box motif. Competitors were the same fragments but were non-labeled. MBP-ERF4 and GST-MYB52 fusion proteins were purified. **(E)** Dual-LUC transient transcriptional activity assays were performed in wild-type Arabidopsis protoplast. Effector and reporter plasmids were prepared as mentioned in Figure 3F with *35S::MYB52* being additionally constructed. Expression of *REN* was used as internal control. LUC/REN represents the relative activity of promoters from different co-transformations. Three independent experiments were performed. The values present the means ± SD. n. s., not significant; **, *P* < 0.01; ***, *P* < 0.001, Student’s t-test.

To test whether the ERF4-MYB52 interaction affects the binding of MYB52 to the *PMEI6*, *14* and *SBT1.7* promoters as well, EMSA assays were performed as described above, but with probes only containing the MBS motif. Similarly, MYB52 alone bound to these MBS motifs (Figure 6A to C, lane 3), while ERF4 did not (Figure 6A to C, lane 2), and the binding of MYB52 to these promoters gradually weakened with the increase in the amount of ERF4 added (Figure 6A to C, lane 4 to 6). Effector plasmids of *proPMEI6*, *14* or *SBT1.7::LUC* were constructed and relative LUC activities were measured. The ERF4-MYB52 interaction decreased the activating regulation of both *PMEI6*, *14* and *SBT1.7* by MYB52, compared to a single co-transformation of *35S::MYB52* with either effector plasmid (Figure 6D).

**Figure 6.**
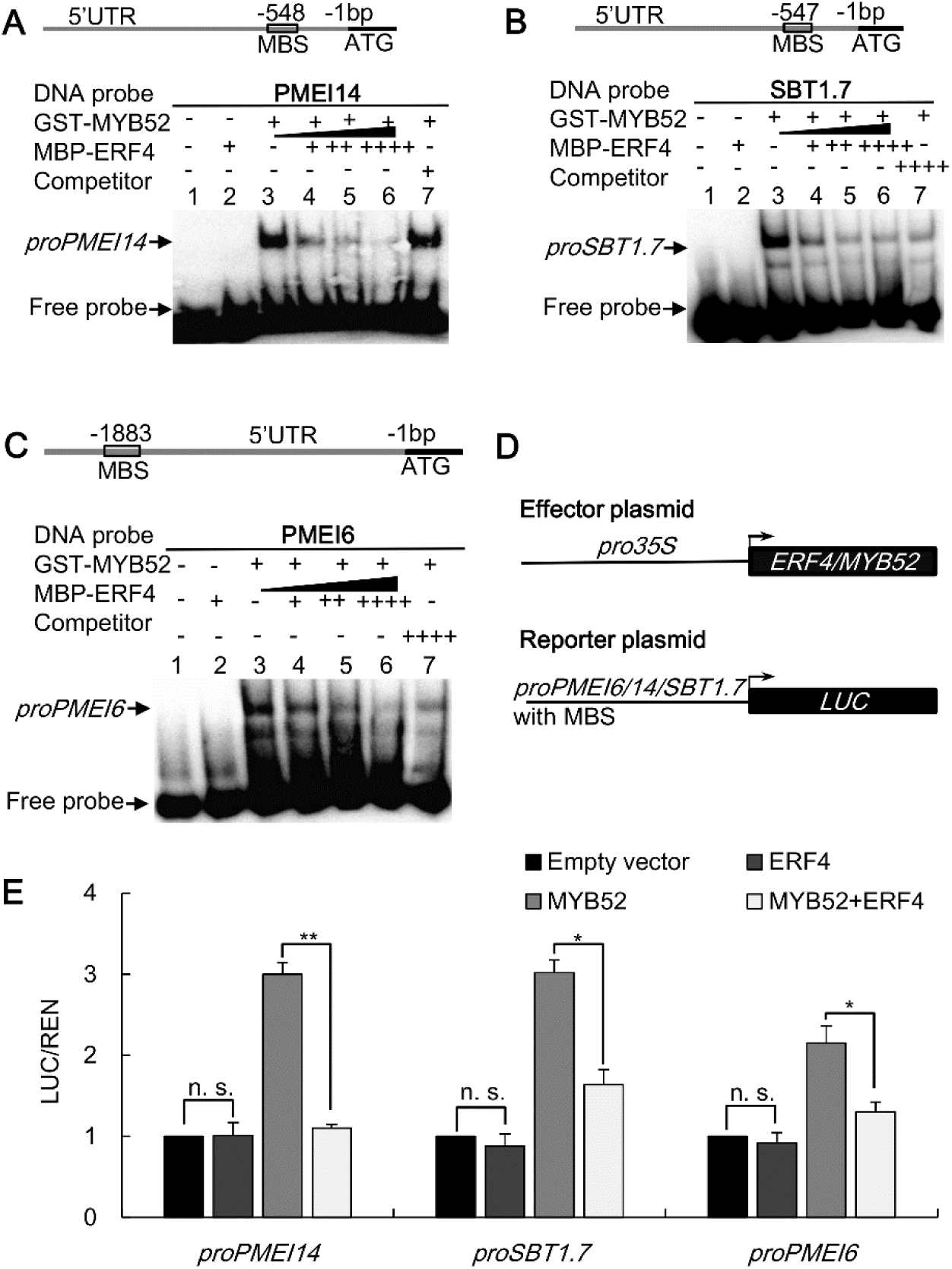
ERF4 interfered MYB52 function in activating downstream genes. **(A)** to **(C)** EMSA results showing that ERF4 did not bind to the MBS motifs in the *PMEI14* **(A)**, *PMEI6* **(B)** and *SBT1.7* **(C)** promoters (lane 2), but MYB52 did bind to them (lane 3). With the increasing of ERF4, the binding of MYB52 to these promoters weakened (lane 4 to 6). Relative nucleotide positions of MBS are indicated (with the first base preceding the ATG start codon being assessed as -1). The DNA probes were 5’biotin-labeled fragments containing the MBS motif. Competitors were the same fragments but were non-labeled. MBS, MYB binding site. **(D)** to **(E)** LUC/REN showing that the activation of MYB52 to the *PMEI6*, *PMEI14* and *SBT1.7* promoters was inhibited by ERF4. The values represent means of three biological replicates ± SD. n. s., not significant; *, *P* < 0.05; **, *P* < 0.01, Student’s t-test.

To further dissect the relationship between ERF4 and MYB52 in the regulation of pectin de-methylesterification, we crossed *erf4-2* and *myb52* and examined the mucilage phenotype of the *erf4-2 myb52* double mutant seeds under vigorous shaking conditions. We observed that disruption of MYB52 function restored the mucilage phenotype of the *erf4-2* mutant, and vice versa (Figure 7A and B). We also examined the expression of *PMEI6*, *13*, *14*, *15* and *SBT1.7* in developing seeds of *erf4-2 myb52*, finding that the expression of *PMEI13*, *14*, *15* was increased whereas *PMEI6* and *SBT1.7* were down-regulated compared to wild-type. However, they all presented a compromised expression pattern compared to that of *erf4-2* and *myb52*, respectively (Figure 2G). Consequently, PME activity of *erf4-2 myb52* and wild-type seeds was alike (Figure 7C), suggesting a complementary effect on DM in the *erf4-2 myb52* seed mucilage. Taken together, these results suggest that ERF4 and MYB52 play completely opposite roles in the same pathway in regulating pectin de-methylesterification in the seed coat mucilage.

**Figure 7.**
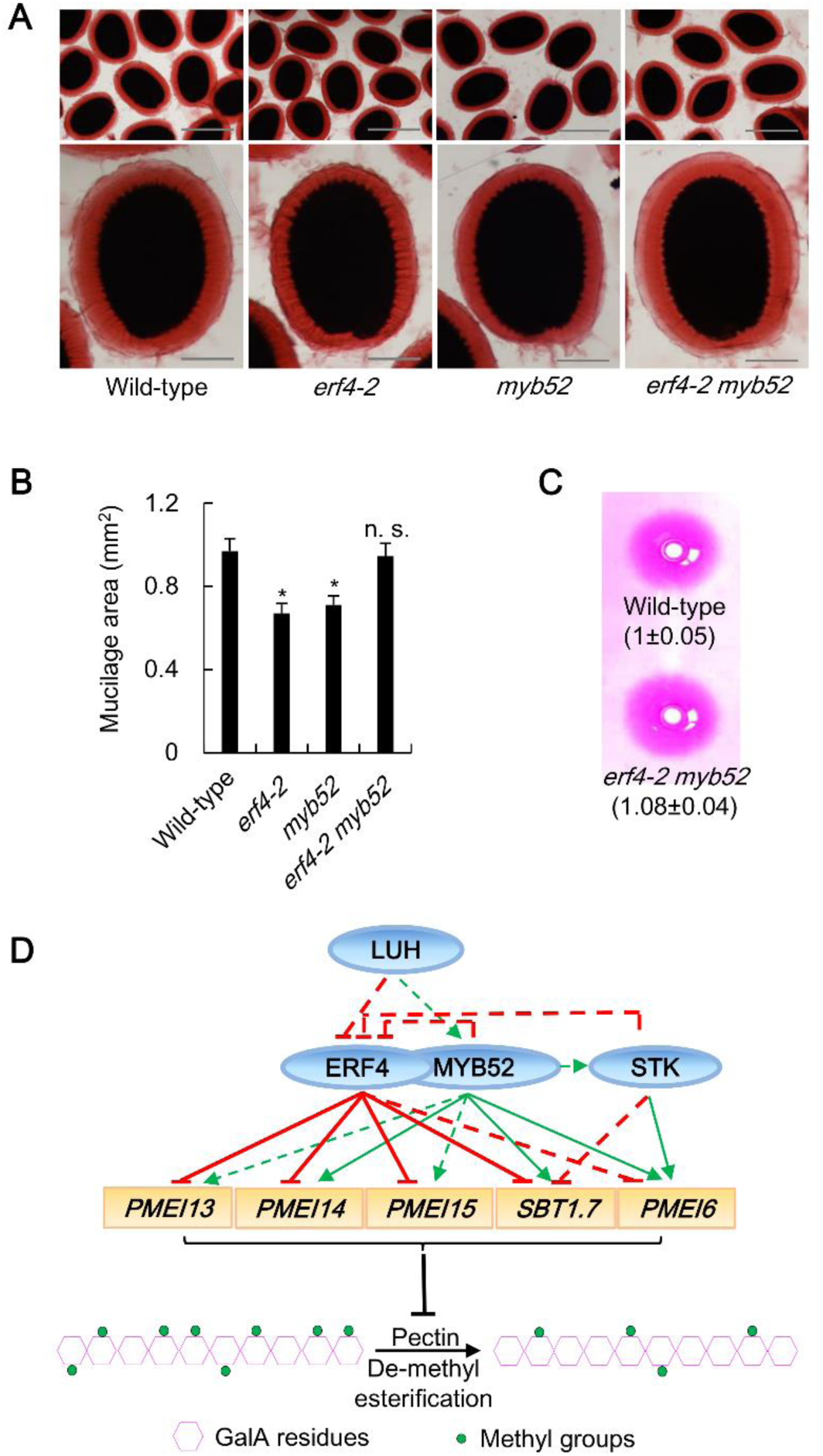
ERF4 and MYB52 antagonize each other’s function in regulating pectin de-methylesterification. **(A)** Mucilage phenotypes for wild-type, *erf4-2*, *myb52* and *erf4-2 myb52* double mutants. Mature dry seeds were shaking in distilled water with 0.01% RR at 200 rpm for 1 h. Bars = 500 μm and 100 μm, respectively. **(B)** Area of adherent mucilage layers. The presented values correspond to average area of AM layer ± SD of at least 20 seeds. n. s., not significant; *, *P* < 0.05, Student’s t-test. **(C)** PME activity of total protein extracts from whole seeds of wild-type and *erf4-2 myb52* double mutants in gel diffusion assay. The values were measured according to gel diffusion and were normalized to average wild-type activity (=1). Errors represent SD values of three biological experiments. **(D)** A model showing that pectin de-methylesterification is under a fine-tuned regulation of the ERF4 and MYB52 transcription complex in the seed coat mucilage. ERF4 and MYB52 interact and suppress each other’s function in binding down-stream genes encoding PME inhibitors and a protease, SBT1.7. ERF4 promotes pectin de-methylesterification by directly suppressing *PMEI13*, *PMEI14*, *PMEI15* and *SBT1.7*, and indirectly suppressing *PMEI6* expression by preventing MYB52 from binding to its promoter. MYB52 negatively regulate pectin de-methylesterification by direct transcriptional activating *PMEI6*, *PMEI14* and *SBT1.7*, and by indirect activating *PMEI13* and *PMEI15* through inhibiting the binding of ERF4 to their promoters. *ERF4* is also negatively regulated by LUH, STK and MYB52 transcription factors. Taken together, ERF4 and MYB52 play completely opposite but precise roles in regulation pectin de-methylesterification in the seed mucilage. The regulatory network found in this study was marked in red, while results from published data in green.

## DISCUSSION

### *ERF4* plays a regulatory role in seed coat mucilage development

The seed coat epidermal differentiation starts after fertilization and proceeds until the mature green stage, corresponding to 0~13 DPA (Francoz et al., 2015; Golz et al., 2018). The *ERF4* expression is gradually increased during this differentiation phase, and peaks at 13 DPA (Figure 1A). At this point in seed development, mucilage synthesis is complete and structural modifications may occur (Francoz et al., 2015; Hu et al., 2016). Since its expression in pod is barely detectable, the detected *ERF4* transcripts are mainly from developing seeds. *ERF4* expression in the seed coat has been demonstrated in a global gene expression dataset (Le et al., 2010). We also observed that it is indeed strongly expressed in epidermal cells of the seed coat, which can be observed with GUS staining (Supplemental Figure 2J and K).

Although *ERF4* has diverse functions in plant growth and development (Koyama et al., 2013; Liu et al., 2017; McGrath et al., 2005; Zhou et al., 2016), its role in cell wall organization has not yet been reported. The spatio-temporal expression patterns of *ERF4* were in agreement with a possible role in mucilage polysaccharides organization during seed development. As expected, we found a modified distribution of the seed mucilage, from adherent to soluble, in the *erf4* mutants (Figure 1E and F; Supplemental Table 1), which is the result of reduced PME activity (Figure 2E and F) along with increased HG methylesterification level (Figure 2A). As the DM of HG domains is supposed to play significant roles in mucilage RG-I solubility (Saez-Aguayo et al., 2013), these results therefore indicate that *ERF4* is involved in post-deposition modification of the seed mucilage polysaccharides by modulating PME activity.

### ERF4 positively modulates HG de-methylesterification by suppressing *PMEI13*, *14*, *15* and *SBT1.7*

Pectin is synthesized in the Golgi apparatus with HGs in a fully methylesterified status (Pelloux et al., 2007; Turbant et al., 2016). The de-methylesterification by PMEs is considered to facilitate the formation of the “egg-box” structures between HG molecules and Ca^2+^, which is deemed to be associated with an increase in cell wall rigidity (Micheli, 2001; Pelloux et al., 2007; Willats et al., 2001a; Willats et al., 2001b; Wolf et al., 2009). The DM in the *erf4* seed mucilage was increased by 10~20% compared to wild-type (Figure 2A), which was confirmed in our immunolabeling assays with increased JIM7 labelling in contrast to weakened JIM5 and CCRC-M38 signals (Figure 2B to D). These results indicate that HG de-methylesterification was inhibited because of the *ERF4* mutation, which might lead to a theoretical decrease in the formation of the “egg-box” structures as verified by 2F4 labelling (Supplemental Figure 5). Thus, at first sight, high level of DM due to the *ERF4* mutation might be associated with a decrease in cell wall rigidity, with consequences on the low cohesiveness of adherent mucilage to the seed coat. Pre-treatment of *pmei6*, *sbt1.7*, *luh/mum1*, *stk* and *myb52* seeds with EDTA, a Ca^2+^ chelator, can promote mucilage extrusion or repartition, because EDTA can destroy the calcium bridge and thus decrease adherence of the inner layer to the seed coat. However, pre-treatment of *erf4* seeds with EDTA has no effects on mucilage partitioning compared to wild-type (Figure 1G; Supplemental Figure 3H). These results indicate that the lost outer part of the *erf4* adherent mucilage in water is insensitive to EDTA extraction.

We proved that the elevated DM was due to a decrease in PME activity (Figure 2E and F). Considering that ERF4 is a transcription repressor (Yang et al., 2005), the reduced PME activity might result from direct up-regulation of rival genes, such as *PMEIs*. We showed that ERF4 suppresses the expression of *PMEI13*, *14*, *15* and *SBT1.7* by directly binding to their regulatory elements that locate in the promoter, intron or exon both *in vitro* and *in vivo* (Figure 3). In most cases, transcription factors regulate gene expression by binding *cis*-acting elements that locate in its promoter or intron. However, *cis*-elements located in exon region were also reported (de Vooght et al., 2008). As *PMEI13*, *14*, *15* and *SBT1.7* are all inhibitors of PME activity, we concluded that the reduced PME activity is due to transcriptional up-regulation of these genes in the *erf4* seeds. Genetic evidence that *erf4-2 pmei13-1* and *erf4-2 pmei14* double mutants showed partially rescued mucilage phenotype further proved that ERF4 functions upstream of *PMEI13*, *14* (Supplemental Figure 9). Attributing to the enormous effect of the *SBT1.7* in DM modification, the *erf4-2 sbt1.7* double mutant seeds resemble *sbt1.7* in mucilage extrusion. We can also conjecture that *SBT1.7* is not primarily regulated by ERF4. Moreover, we cannot exclude the contributions of other *PMEI* genes to the *erf4* mucilage phenotype (such as *PMEI6*, *AT1G11593*, etc.), since their expressions were also up-regulated in the *erf4* seeds (Supplemental Figure 7A).

*PMEI6*, *14*, *PME58*, *FLY1* and *SBT1.7* have been proved to be involved in HG de-methylesterification in the seed coat mucilage (Rautengarten et al., 2008; Saez-Aguayo et al., 2013; Shi et al., 2018; Turbant et al., 2016; Voiniciuc et al., 2013). In this study, we showed that *PMEI13* and *PMEI15* play a role in HG de-methylesterification in the seed mucilage, although *PMEI15* might play only a supporting role (Supplemental Figure 9). The transcriptional regulation network of HG de-methylesterification is also partially elaborated, although it is just the tip of the iceberg. LUH/MUM1, STK and MYB52, either positive or negative regulator of mucilage pectin methylesterification modification, are transcriptionally associated with each other in regulating DM related genes (Shi et al., 2018). In this study, we showed that ERF4 functions in regulating HG de-methylesterification by suppressing *PMEI13*, *14*, *15* and *SBT1.7*. In addition, *ERF4* expression was suppressed by MYB52, STK and LUH/MUM1 in our qRT-PCR analysis (Figure 2G). The above results suggest that during seed mucilage maturation, the regulatory network underlying pectin de-methylesterification is more complex than expected.

### Possible influences of PME activity on seed development

Plant cell growth is regulated by the interplay between the intracellular turgor pressure and the flexibility of the cell wall. Many plant cell wall-modifying enzymes, such as PMEs and PME inhibitors play important roles in cell wall reorganization for control of a variety of growth processes (Levesque-Tremblay et al. 2015b), including fruit ripening (Reca et al., 2012), cell elongation (Derbyshire et al., 2007; Guenin et al., 2011), stress responses (Bethke et al., 2014; Lionetti et al., 2007) and mediating unilateral cross-incompatibility (Zhang et al., 2018).

The functional roles of pectin methylesterification in regulating seed growth and seed size have also been reported. For example, Arabidopsis plants overexpressing *PMEI5* showed a significant enlargement of seed size (Muller et al., 2013). *stk* seeds presented a reduced seed size phenotype accompanied with elevated PME activity (Ezquer et al., 2016). These findings indicate that pectin de-methylesterification is negatively correlated with seed growth. Conversely, mutants of PME encoding genes, *hms* and *pme58*, displayed a reduced cell size in the embryo and seed epidermal cells which is a consequence of lack of cell expansion (Levesque-Tremblay et al., 2015a; Turbant et al., 2016). In this study, we found that both seed size and mucilage-producing cell surface area of *35S::ERF4* were increased compared to wild-type seeds (Supplemental Figure 6). Collectively, higher DM might be correlated with smaller seed size in the cases of *35S::ERF4*, *hms* and *pme58* mutants, but not in *stk* and *35S::PMEI5* seeds.

Previous studies have established a complex relationship between DM and plant growth processes. It has been demonstrated that higher level of DM seems to be able to either promote or inhibit cell expansion and organ growth (Wormit and Usadel, 2018). For instances, overexpressing *AtPMEI2* resulted in enhanced root elongation, whereas *AtPMEI4* and *OsPMEI28* overexpressors have delayed hypocotyl growth rate and restrained culm elongation, respectively (Nguyen et al., 2016; Pelletier et al., 2010; Rockel et al., 2008). Opposed to *AtPME1* which inhibits pollen tube growth, Arabidopsis *VANGUARD1* is required for the growth of pollen tubes (Jiang et al., 2005). Pectin de-methylesterification manipulated by *AtPME5* also contributes to an increase in elasticity of the shoot apical meristem (Peaucelle et al., 2011). These findings suggest that PME activity could be tightly controlled to fine-tune pectin’s biophysical properties in the cell walls of certain organs, resulting in different modes of action of DM in the development of different organs. In summary, we conjectured that pectin de-methylesterification catalyzed by PMEs could either promote or inhibit cell expansion in developing seeds, which might be correlated with the modification patterns/degree of the cell wall. Future studies are needed to elucidate the correlations between these patterns resulting from cell wall-modifying enzymes and the flexibility of the plant cell wall structure, and how they affect cell expansion.

### ERF4 and MYB52 antagonistically regulate pectin de-methylesterification in the seed mucilage

In eukaryotes, transcription factors usually work in combination which can promote or inhibit each other’s transcriptional activity in controlling expression of target genes. For example, the BRC1 transcriptional activity is suppressed by interaction with the transcriptional repressor TIE1 in Arabidopsis (Yang et al., 2018); the interaction between MdMYC2 and MdERF2 not only inhibits the binding of MdERF2 to *MdACS1*, but also prevents MdMYC2 from binding to *MdACS1* and *MdACO1* in apple (Li et al., 2017); and during crown root elongation, the WOX11-ERF3 interaction either enhances WOX11-mediated repression or inhibits ERF3-mediated activation of *RR2* in rice (Zhao et al., 2015). Here, we provided evidence for an interaction between ERF4 and MYB52 (Figure 4). One effect of this interaction might be on the regulation of ERF4 to its downstream genes. We showed that the presence of MYB52 inhibits the binding of ERF4 to *PMEI13*, *14*, *15* and *SBT1.7* and enhanced their transcription activity (Figure 5), indicating that the ERF4-MYB52 interaction represses the transcriptional suppression of ERF4 to *PMEI13, 14, 15* and *SBT1.7*, thereby decreasing PME activity in the seed coat mucilage. We also observed that the ERF4-MYB52 interaction prevents the binding of MYB52 to *PMEI6, 14* and *SBT1.7* promoters, and reduces its activating regulation (Figure 6), suggesting that this interaction suppresses the binding of MYB52 to the *PMEI6*, *14* and *SBT1.7* promoters, thereby enhancing PME activity in the seed mucilage.

From these downstream targets of the ERF4-MYB52 transcriptional complex, *PMEI13* and *PMEI15* have GCC-box as well as MBS motifs in their promoters, but MYB52 did not recognize these MBS motifs in our study by both EMSA and ChIP assays (data not shown), whereas *PMEI6* only has one MBS in its promoter. However, in our qRT-PCR experiments, the expression of *PMEI6* was up-regulated in *erf4-2*, and the *PMEI13*, *15* transcripts were decreased in *myb52* (Figure 2G). We reasoned that another effect of the ERF4-MYB52 interaction might be the indirect transcriptional regulation of these genes, that MYB52 promotes *PMEI13* and *PMEI15* expression by inhibiting the transcriptional suppression activity of ERF4, and ERF4 suppresses *PMEI6* expression by antagonizing the function of MYB52.

The reciprocal transcriptional inhibition between ERF4 and MYB52 provides insights into the mechanisms that plants use to maintain the appropriate DM, to further regulate plasticity of the cell wall structure in the seed mucilage. It was further verified by genetic evidence that PME activity and mucilage release of the *erf4-2 myb52* double mutant are almost identical to those of wild-type (Figure 7A to C). Moreover, according to the expression patterns of downstream genes in *erf4-2*, *myb52* and *erf4-2 myb52* (Figure 2G), we conjectured that for the ERF4-MYB52 transcriptional complex *PMEI13, 14, 15* could be mainly regulated by ERF4 whereas *PMEI6* and *SBT1.7* could be mainly under positive regulation of MYB52, although other transcription factors might also be involved. A model for the role of the ERF4-MYB52 complex in the regulation of pectin de-methylesterification was proposed (Figure 7D): (1) ERF4 and MYB52 interact and suppress each other’s transcriptional activity; (2) ERF4 negatively regulates *PMEI13, 14, 15* and *SBT1.7* expression by directly binding to their regulatory elements and suppresses *PMEI6* indirectly by antagonizing MYB52 function, giving rise to positive regulation of pectin de-methylesterification; (3) MYB52 activates *PMEI6*, *14* and *SBT1.7* by directly binding to their promoters and positively regulates *PMEI13*, *15* expression indirectly by suppressing ERF4 activity, which in turn negatively regulates pectin de-methylesterification in the seed mucilage. Taken together, these findings show a compensatory effect on overall degree of HG methylesterification, where ERF4 and MYB52 antagonistically modulate the structure of adherent mucilage by regulating genes involved in HG de-methylesterification in the seed coat mucilage.

## METHODS

### Plant materials and growth conditions

The Arabidopsis T-DNA insertion lines *erf4-1* (SALK_200761), *erf4-2* (SALK_073394), *myb52* (SALK_138624), *pmei13-1* (GABI-601A06), *pmei13-2* (SALK_038767), *pmei14* (SALK_206157), *pmei15-1* (SALK_106719), *pmei15-2* (SALK_053746) and *sbt1.7* (GABI-140B02) were obtained from Nottingham Arabidopsis Stock Centre (http://arabidopsis.info/). The *myb52*, *pmei14* and *sbt1.7* mutants have been described previously (Rautengarten et al., 2008; Shi et al., 2018). The Arabidopsis wild-type (Col-0 ecotype) and mutant plants were grown at 22 °C under long-day (16 h light/8 h dark) conditions in a growth chamber. Double mutant was obtained by crossing *erf4-2* with either *myb52*, *pmei13-1*, *pmei14*, *pmei15-1* or *sbt1.7* single mutant. In all comparative analysis for mucilage extrusion phenotype, seeds used from mutants and wild-type plants were simultaneously cultivated and harvested.

### PCR-based genotyping

Homozygous T-DNA insertion lines for all the single and double mutants were identified by PCR using primers (Supplemental Table 2) provided by T-DNA Primer Design (http://signal.salk.edu/tdnaprimers.2.html) with genomic DNA extracts. Plants with PCR products only obtained for the insertion border and not with primers flanking the insertion sites were regarded as homozygous lines.

### Expression analysis and GUS staining of plant tissues

Total RNAs were extracted from 4 to 13 DPA developing seeds of wild-type and mutants including *erf4-2*, *myb52*, *luh*, *stk*, *pmei13*, *pmei15*, *35S::ERF4* and *erf4-2 myb52*, as well as Arabidopsis organs such as roots, seedlings, leaves and inflorescence stems using the RNeasy plant mini kit (Qiagen). Total amount of 1 μg RNA was used as template for first-strand cDNA synthesis with the oligo(dT) primer and the PrimeScript^TM^RT reagent Kit with gDNA Eraser (TaKaRa) according to the manufacturer’s instructions. The qRT-PCR assay was performed using 1/20 diluted cDNA as templates in the reactions containing SYBR^®^ Premix Ex Taq^TM^ II (TaKaRa). The qRT-PCR assay was conducted in triplicate in an ABI 7500 Fast Real-Time PCR System. The relative expression of genes was calculated by normalizing against a housekeeping gene *ACTIN2* (Dekkers et al., 2012) with the 2^−ΔΔC^_T_ method in analyzing the data. *ACTIN2* was also used as loading control in the RT-PCR analysis for detecting of transcripts.

The *ERF4* promoter region of 1,965 bp preceding the transcriptional start codon ATG was amplified by PCR using wild-type genomic DNA as template. After purification with the EasyPure^®^ PCR Purification Kit (TRANSGEN BIOTECH), the PCR products were recombined into the PBI121 binary vector with *XbaI* site to generate *proERF4::GUS*. After *Agrobacterium*-mediated transformation of the Arabidopsis wild-type plants, histochemical staining of positive transformants was performed using the GUS staining kit (Solarbio) according to the manufacturer’s instructions. The GUS stained tissues were examined using a stereoscopic microscope (LEICA MDG29) equipped with a Leica MC190 camera. Primers used are listed in Supplemental Table 2.

### Ruthenium red staining and morphological analysis

For mucilage extrusion analysis, whole mature dry seeds were shaking by hand or in an incubator (under 28 °C, 200 rpm,1 h conditions) in deionized water, 50 mM EDTA (pH 8.0) or 0.01% (w/v) ruthenium red (RR, Sigma-Aldrich). After RR staining, seeds were rinsed in deionized water and visualized with a bright-field microscope (Nikon). The seed size and area of mucilage halos were measured with the Image J software (https://imagej.en.softonic.com/download). Dry seeds were mounted on stubs using an adhesive disc and then coated with platinum using a Hitachi E1045 ion sputter. The morphology of MSCs surface of mature dry seeds was investigated by scanning electron microscopy (SEM, Hitachi S4800) at an accelerating voltage of 20 kV. Measurement of MSC surface areas were carried out with the Image J software, and more than 300 measurements were performed. For the morphological analysis of seed coat differentiation, developing seeds at stages 7 and 10 DPA of wild-type and *erf4* were sectioned and stained with 0.5% (w/v) Toluidine Blue O (TBO) after fixing and embedding with spurr resin as described by Shi et al. (2018). The sections were visualized and photographed with a bright-field microscope (Nikon).

### Gene cloning and plant transformation

The full-length CDS of *ERF4* was amplified by PCR using wild-type genomic DNA as template and was recombined into *p*CAMBIA35tlegfps2#4 binary vector with *KpnI/XbaI* restriction sites to generate *35S::ERF4* overexpressing vector. The full-length CDS of *ERF4* was also recombined into a modified *p*CAMBIA1300 binary vector (*35S::Myc*) with the *Kpn I* site to obtain *35S::Myc-ERF4* overexpression vector. The vectors were then introduced into the *Agrobacterium* strain GV3101 to transform the Arabidopsis wild-type or *erf4-2* mutant plants by a floral dip method to obtain overexpression and complementary lines. Positive transformants were selected on 1/2 MS medium containing 25 mg L^−1^ hygromycin or 50 mg L^−1^ kanamycin.

### Mucilage extraction and monosaccharide composition analysis

Briefly, 5 mg mature dry seeds from wild-type or mutants with three biological replicates were placed in 2 mL tubes with 1 mL deionized water. The water-soluble mucilage was extracted in an incubator under 28 °C and 200 rpm (1 h) conditions. The seeds were washed three times, and supernatants were pooled in 10 mL glass tubes. The seeds were then treated with ultrasonic for 20 s in 1 mL deionized water to obtain adherent mucilage as described by Zhao et al., (2016). The de-mucilaged seeds were washed three times, and pooled supernatants were placed in 10 mL glass tubes. Non-adherent mucilage was also solubilized with 50 mM EDTA (pH 8.0) using the same procedure described for water extraction, whereas adherent mucilage was extracted by shaking seeds in EDTA with the TissueLyser II (Qiagen) at 20 movements/s for 20 min. The mucilage extractions were then lyophilized with a FreeZone® freeze dryer (LABCONCO), which were used for compositional monosaccharide analysis afterwards. Pictures of RR-stained seeds post-extraction are shown in Supplemental Figure 2. To determine monosaccharide composition of both fractions, the mucilage polymers were hydrolyzed to monose with 2 M trifluoroacetic acid (110 °C, 2 h) and quantified by HPLC (Waters 2695 and 2998) after 1-phenyl-3-methyl-5-pyrazolone derivatization (70 °C, 0.5 h) as described by Shi et al. (2018).

### Determination of PME activity and degree of pectin methylesterification

Gel diffusion assays were performed to determine PME activity. In brief, 20 mg mature dry seeds were grounded into homogenate with mortar and pestle in 400 µL of extraction buffer (1 M NaCl, 12.5 mM citric acid, and 50 mM Na_2_HPO_4_, pH 6.5) to obtain total protein extracts. The resulting homogenate was centrifuged at 20,000 *g* for 15 min after shaking at 4 °C for 1h. Protein concentrations in the supernatant were determined according to the Bradford method (Bradford, 1976). Equal quantities of proteins (10 µg) in the same volume (20 µL) were loaded into 6-mm-diameter wells in 1% agarose gels containing 0.1% (w/v) of esterified citrus fruit pectin (85% esterified, Sigma-Aldrich), 12.5 mM citric acid and 50 mM Na_2_HPO_4_, pH 6.5. After incubation overnight at 28 °C, the gels were stained for 45 min with 0.01% RR and washed five times in 5 h with water. The gels were photographed and the red-stained areas were quantified with the Image J software. The measurements were performed in triplicate and relative PME activity was normalized with the wild-type average area being set to 100%.

Water-extracted whole mucilage from 20 mg seeds by ultrasonic were used to determine DM according to Voiniciuc et al. (2013). In brief, methanol was released from mucilage by alkaline de-esterification with 2 M NaOH at 4 °C for 1 h. After neutralization of extracts with 2 M HCl, released methanol was oxidized with alcohol oxidase (0.5 U, Sigma-Aldrich) at 25 °C for 15 min. Thereafter, a mixture containing 20 mM 2, 4-pentanedione in 2 M ammonium acetate and 50 mM acetic acid was added. After incubation at 60 °C for 15 min, samples were directly cooled on ice. Absorbance at 412 nm was measured with a plate reader (Tecan). The methanol content was calculated as the amount of formaldehyde produced from methanol by alcohol oxidase, by comparison with a standard calibration curve (Klavons and Bennett, 1986). Meanwhile, add 220 µL sodium tetraborate/sulfuric acid into 40 µL supernatant from the remaining mucilage saponification solution on ice before incubation at 100°C for 5 min. After being directly cooled on ice, samples are mixed with 4 µL 1.5 mg/mL M-hydroxybiphenyl. Absorbance was measured at 525 nm with a plate reader (Tecan). GalA was quantified using a D-(+)-galacturonic acid monohydrate (Sigma-Aldrich) standard curve. DM = total methanol content (µmol)/total GalA content (mg) × 100%.

### Immunolabeling assay

Monoclonal antibodies CCRC-M38 (specifically recognize de-esterified HG), JIM5 (specifically recognize low methylesterified HG), JIM7 (specifically recognize fully methylesterified HG) and 2F4 (specifically recognize Ca^2+^-linked HG dimers) were used for whole-seed immunolabelling analysis. CCRC-M38, JIM5 and JIM7 antibodies were used with PBS buffer (140 mM NaCl, 2.7 mM KCl, 8.0 mM Na_2_HPO_4_, 1.5 mM KH_2_PO_4_, pH7.4), whereas 2F4 antibody required TCS buffer (20 mM Tris-HCl pH 8.2, 0.5 mM CaCl_2_, 150 mM NaCl). Whole intact dry seeds were firstly blocked in 3% (w/v) fat-free milk powder in PBS/TCS buffer (MPBS/MTCS) at 37 °C for 1 h, and then labeled with 10-fold MPBS/MTCS-diluted primary antibody at 37 °C for 1.5 h. Seeds were subsequently washed 3 times in PBS/TCS buffer. Then 200-fold MPBS/MTCS-diluted secondary antibody AlexaFluor488-tagged donkey anti-rat IgG (Thermofisher) antibody for JIM5 and JIM7, and a donkey anti-mouse IgG (Thermofisher) for CCRC-M38 and 2F4 were used to incubate with seeds at 37 °C for 1.5 h in dark. Finally, seeds were double labeled with Calcofluor White (Sigma-Aldrich) which was diluted 5 times in PBS/TCS buffer for 15 min. Images were captured by using a FluoView FV1000 spectral confocal laser microscope (OLYMPUS) with 405 and 488 nm laser.

### EMSA assay

The induction and purification of proteins were described as (Yuan et al., 2014). In brief, The CDS of MYB52 was cloned into *p*GEX4T-1 vector to generate GST-MYB52 fusion protein, whereas the CDS of ERF4 was cloned into *p*MAL-C2X vector in order to generate MBP-ERF4 fusion protein. The resulting plasmids were separately transformed into *Escherichia coli* strain BL21 for induction of fusion proteins. Empty *p*GEX4T-1 and *p*MAL-C2X vectors were also transformed to obtain GST and MBP tags for control experiments. The concentration of isopropyl β-D-thiogalactoside (IPTG) for protein induction was 0.5 mM. The purification of GST tag or GST-MYB52 fusion protein was performed using a GST-tag Protein Purification Kit (Beyotime, P2262) with the BeyoGold™ GST-tag Purification Resin according to the user guide. The purification of MBP tag or MBP-ERF4 fusion protein was performed with the PurKine MBP-Tag Protein Purification Kit (Dextrin) as recommended by the manufacturer. Oligonucleotide probes were synthesized with their 5’-end being labelled with biotin. To prepare double strand probes, forward and reverse oligonucleotide probes were heated at 95 °C for 5 min in annealing buffer, and then naturally cooled to room temperature. The EMSA assay was performed using a LightShift™ Chemiluminescent EMSA Kit (ThermoFisher) according to instructions recommended by the manufacturer. All primers and oligonucleotide probes used were listed in Supplemental Table 2.

### ChIP-qPCR assay

About 2 g of immature siliques at 10-13 DPA stage were collected from wild-type and *pro35S::Myc-ERF4* or *pro35S::Myc-MYB52* transgenic plants with three repeats. After washing two times with ddH_2_O, siliques were fixed in 37 mL 1% formaldehyde in vacuum for 10 min. After termination of the crosslinking with 2.5 mL 2 M Glycine, samples were ground to a fine powder in liquid nitrogen. Immunoprecipitation of chromatin was performed as previously described (Gendrel et al., 2005) with anti-Myc antibody (Abcam). The precipitated DNA was recovered and the enrichment of DNA fragments in the immunoprecipitated chromatin were quantified by qPCR analysis using primers listed in Supplemental Table 2.

### Dual-luciferase transient transcriptional activity assay

The modified *p*BI221 vector which removed the *GUS* reporter gene between the *Kpn*I and *Bam*HI sites was used to create the effector constructs of *pro35S::ERF4* and *pro35S::MYB52* with the CDS of *ERF4* and *MYB52*. To generate *proPMEI13::LUC*, *proPMEI14i::LUC*, *proPMEI15::LUC*, *proSBT1.7e::LUC*, *proPMEI14::LUC*, *proPMEI6::LUC* and *proSBT1.7::LUC* reporter plasmids, genomic DNA sequences containing the GCC-box or MBS motifs of the *PMEI13* promoter (2,652 bp), *PMEI14* intron (1,575 bp) and promoter (1,047 bp), *PMEI15* promoter (1,188 bp), *SBT1.7* exon (1,186 bp) and promoter (893 bp) and *PMEI6* promoter (1,998 bp) were cloned into *Hind* III and *Bam*HI sites of the *p*GreenII-0800 vector. The empty vector was used as negative control. Dual-luciferase transient assays were performed in Arabidopsis mesophyll protoplasts from 4-week old plants. The ratio between LUC and REN activity was measured with three biological replicates.

### GST pull-down assay

Pull-down experiments were performed with the purified GST-MYB52 and MBP-ERF4 fusion proteins. Purified GST and MBP tag were used as negative controls. The purified GST tag or GST-MYB52 fusion protein were used as bait after removing of reduced glutathione. After binding of GST tag or GST-MYB52 to Glutathione Sepharose 4B resin, purified MBP tag or MBP-ERF4 fusion protein were additionally added to the column and incubated at 4 °C for 1 h on a rotator, then unbound proteins were washed away with the Binding Buffer. The bound proteins were fractionated on 10% SDS-PAGE gel after boiled in water for 5 min. Western blot was performed as described (Yuan et al., 2014) with an anti-MBP antibody (Abcam).

### Co-IP assay

The co-IP experiments were performed as described (Yao et al., 2017). Briefly, The CDS of *MYB52* and *ERF4* were individually introduced into *p*CAMBIA1307-Myc and *p*CAMBIA1307-Flag vectors to generate *pro35S::MYB52-Myc* and *pro35S::ERF4-Flag*. Different combinations of plasmids were transformed into the Arabidopsis mesophyll protoplasts. The transformed cells were cultured at 22 °C for 16 h, prior to harvest for protein extraction. The anti-Myc antibody (Abcam) and protein A+G agarose beads were used to precipitate the MYB52-Myc and ERF4-Flag complex. Proteins were fractionated on 10% SDS-PAGE gel after boiled in water for 5 min for use in immunoblot analysis with the anti-Flag antibody (Sigma-Aldrich).

### BiFC assay

The BiFC experiments were performed as described (Wang et al., 2019). Briefly, the CDS of *ERF4* was fused with the C-terminal of the yellow fluorescent protein (YFP) in *p*E3242 vector to generate *pro35S::ERF4-YC*, and that of *MYB52* was fused with the N-terminal of YFP in *p*E3228 vector in order to generate *pro35S::YN-MYB52*. The constructed plasmids were transformed into Arabidopsis mesophyll protoplasts in different combinations. The transfected cells were incubated in dark for 12 to 16 h. Images were captured by using a FluoView FV1000 spectral confocal laser microscope (OLYMPUS). The YFP was observed with the 514 nm laser. DAPI was observed with the 358 nm laser. Chloroplast (CHl) auto-fluorescence was observed with the 488 nm laser.

## Accession Numbers

Sequence data used in this study can be found in The Arabidopsis Information Resource (TAIR; https://www.arabidopsis.org) under the following accession numbers: ERF4 (At3g15210), MYB52 (At1g17950), LUH/MUM1 (At2g32700), STK (At4g09960), PMEI6 (At2g47670), PMEI13 (At4g15750), PMEI14 (At1g56100), PMEI15 (At3g05741), SBT1.7 (At5g67360) and ACTIN2 (At3g18780).

## Supplemental Data

**Supplemental Figure 1.** Expression pattern of *ERF4, PMEI13, PMEI14, PMEI15* and *SBT1.7* during Arabidopsis plant development.

**Supplemental Figure 2.** GUS activity analysis of the *ERF4* promoter.

**Supplemental Figure 3.** Seed coat morphology analysis and RR stained mucilage halos after water and EDTA extraction.

**Supplemental Figure 4.** Complementation of *erf4-2* with the *35S::ERF4* construct.

**Supplemental Figure 5.** A decreased 2F4 labelling was detected for *erf4-2* seeds compared to that of wild-type.

**Supplemental Figure 6.** Phenotypes of the *35S::ERF4* overexpressors.

**Supplemental Figure 7.** Relative expression of DM related genes in *erf4-2* and *35S::ERF4* overexpressors compared to wild-type.

**Supplemental Figure 8.** The transcriptional suppression of ERF4 to *SBT1.7* was abolished by point mutation in the GCC-box motif that locates in its exon.

**Supplemental Figure 9.** Roles of *PMEI13* and *PMEI15* in mucilage maturation and genetic interactions between ERF4 and its downstream targets.

**Supplemental Table 1.** Composition of sequentially extracted mucilage for wild-type and *erf4* seeds.

**Supplemental Table 2.** Primers used for genotype identification.

**Supplemental Table 3.** Primers used for qRT-PCR and RT-PCR analysis.

**Supplemental Table 4.** Primers used for vector construction.

**Supplemental Table 5.** Probes used in EMSA experiments.

## ACKNOWLEDGMENTS

This work was supported by the National Natural Science Foundation of China (project No. 31670302, 31600237 and 31470291), the Agricultural Science and Technology Innovation Program (ASTIP-TRIC02), the National Key Technology R&D Program (2015BAD15B03-05), the Elite Youth Program of CAAS (to Yingzhen Kong) and the Taishan Scholar Program of Shandong (to Gongke Zhou).

## AUTHOR CONTRIBUTIONS

Yingzhen Kong and Anming Ding designed the research. Anming Ding, Xianfeng Tang, Linhe Han, Jianlu Sun, Angyan Ren, Jinhao Sun, Zongchang Xu and Ruibo Hu each performed some of the experiments. All authors analyzed the data. Anming Ding wrote the paper. Gongke Zhou and Yingzhen Kong revised the manuscript.

## References

Arsovski, A.A., Haughn, G.W., and Western, T.L. (2010). Seed coat mucilage cells of Arabidopsis thaliana as a model for plant cell wall research. Plant Signal Behav 5, 796–801.

Bethke, G., Grundman, R.E., Sreekanta, S., Truman, W., Katagiri, F., and Glazebrook, J. (2014). Arabidopsis PECTIN METHYLESTERASEs contribute to immunity against Pseudomonas syringae. Plant Physiol 164, 1093–1107.

Bradford, M.M. (1976). A rapid and sensitive method for the quantitation of microgram quantities of protein utilizing the principle of protein-dye binding. Analytical Biochem 72, 248–254.

de Vooght, K.M., van Wijk, R., and van Solinge, W.W. (2008). GATA-1 binding site in exon1 direct erythroid-specific transcription of PPOX. Gene 409, 83–91.

Dekkers, B.J., Willems, L., Bassel, G.W., van Bolderen-Veldkamp, R.P., Ligterink, W., Hilhorst, H.W., and Bentsink, L. (2012). Identification of reference genes for RT-qPCR expression analysis in Arabidopsis and tomato seeds. Plant Cell Physiol 53, 28–37.

Derbyshire, P., McCann, M.C., and Roberts, K. (2007). Restricted cell elongation in Arabidopsis hypocotyls is associated with a reduced average pectin esterification level. BMC Plant Biol 7, 31.

Ehlers, K., Bhide, A.S., Tekleyohans, D.G., Wittkop, B., Snowdon, R.J., and Becker, A. (2016). The MADS Box Genes ABS, SHP1, and SHP2 Are Essential for the Coordination of Cell Divisions in Ovule and Seed Coat Development and for Endosperm Formation in Arabidopsis thaliana. PLoS One 11, e0165075.

Ezquer, I., Mizzotti, C., Nguema-Ona, E., Gotte, M., Beauzamy, L., Viana, V.E., Dubrulle, N., Costa de Oliveira, A., Caporali, E., Koroney, A.S., Boudaoud, A., Driouich, A., and Colombo, L. (2016). The Developmental Regulator SEEDSTICK Controls Structural and Mechanical Properties of the Arabidopsis Seed Coat. Plant Cell 28, 2478–2492.

Fan, Z.Q., Kuang, J.F., Fu, C.C., Shan, W., Han, Y.C., Xiao, Y.Y., Ye, Y.J., Lu, W.J., Lakshmanan, P., Duan, X.W., and Chen, J.Y. (2016). The Banana Transcriptional Repressor MaDEAR1 Negatively Regulates Cell Wall-Modifying Genes Involved in Fruit Ripening. Front Plant Sci 7, 1021.

Francoz, E., Ranocha, P., Burlat, V., and Dunand, C. (2015). Arabidopsis seed mucilage secretory cells, regulation and dynamics. Trends Plant Sci 20, 515–524.

Fujimoto, S.Y., Ohta, M., Usui, A., Shinshi, H., and Ohme-Takagi, M. (2000). Arabidopsis ethylene-responsive element binding factors act as transcriptional activators or repressors of GCC box-mediated gene expression. Plant Cell 12, 393–404.

Gendrel, A.V., Lippman, Z., Martienssen, R., and Colot, V. (2005) Profiling histone modification patterns in plants using genomic tiling microarray. Nat method 2, 213–218.

Golz, J.F., Allen, P.J., Li, S.F., Parish, R.W., Jayawardana, N.U., Bacic, A., and Doblin, M.S. (2018). Layers of regulation-Insights into the role of transcription factors controlling mucilage production in the Arabidopsis seed coat. Plant Sci 272, 179–192.

Griffiths, J.S., Crepeau, M.J., Ralet, M.C., Seifert, G.J., and North, H.M. (2016). Dissecting Seed Mucilage Adherence Mediated by FEI2 and SOS5. Front Plant Sci 7, 1073.

Guenin, S., Mareck, A., Rayon, C., Lamour, R., Assoumou Ndong, Y., Domon, J.M., Senechal, F., Fournet, F., Jamet, E., Canut, H., Percoco, G., Mouille, G., Rolland, A., Rusterucci, C., Guerineau, F., Wuytswinkel, O.V., Gillet, F., Driouich, A., Lerouge, P., Gutierrez, L., and Pelloux, J. (2011). Identification of pectin methylesterase 3 as a basic pectin methylesterase isoform involved in adventitious rooting in Arabidopsis thaliana. New Phytol 192, 114–126.

Haughn, G.W., and Western, T.L. (2012). Arabidopsis Seed Coat Mucilage is a Specialized Cell Wall that Can be Used as a Model for Genetic Analysis of Plant Cell Wall Structure and Function. Front Plant Sci 3, 64.

Hu, R., Li, J., Wang, X., Zhao, X., Yang, X., Tang, Q., He, G., Zhou, G., and Kong, Y. (2016). Xylan synthesized by Irregular Xylem 14 (IRX14) maintains the structure of seed coat mucilage in Arabidopsis. J Exp Bot 67, 1243–1257.

Huang, J., DeBowles, D., Esfandiari, E., Dean, G., Carpita, N.C., and Haughn, G.W. (2011). The Arabidopsis transcription factor LUH/MUM1 is required for extrusion of seed coat mucilage. Plant Physiol 156, 491–502.

Jiang, L.X., Yang, S.L., Xie, L.F., Puah, C.S., Zhang, X.Q., Yang, W.C., Sundaresan, V., and Ye, D. (2005). VANGUARD1 encodes a pectin methylestrase that enhance pollen tube growth in the Arabidopsis style and transmitting tract. Plant Cell 17, 584–596.

Jofuku, K.D., den Boer, B.G.W., Montagu, M.V., and Okamuro, J.K. (1994). Control of Arabidopsis flower and seed development by the homeotic gene APETALA2. Plant Cell 6, 1211–1225.

Klavons, J.A., and Bennett, R.D. (1986). Determination of methanol using alcohol oxidase and its application to methyl ester content of pectins. Journal of Agricultural and Food Chemistry 34, 597–599.

Klepikova, A.V., Kasianov, A.S., Gerasimov, E.S., Logacheva, M.D., and Penin, A.A. (2016). A high resolution map of the Arabidopsis thaliana developmental transcriptome based on RNA-seq profiling. Plant J 88, 1058–1070.

Koyama, T., Nii, H., Mitsuda, N., Ohta, M., Kitajima, S., Ohme-Takagi, M., and Sato, F. (2013). A regulatory cascade involving class II ETHYLENE RESPONSE FACTOR transcriptional repressors operates in the progression of leaf senescence. Plant Physiol 162, 991–1005.

Le, B.H., Cheng, C., Bui, A.Q., Wagmaister, J.A., Henry, K.F., Pelletier, J., Kwong, L., Belmonte, M., Kirkbride, R., Horvath, S., Drews, G.N., Fischer, R.L., Okamuro, J.K., Harada, J.J., and Goldberg, R.B. (2010). Global analysis of gene activity during Arabidopsis seed development and identification of seed-specific transcription factors. Proc Natl Acad Sci 107, 8063–8070.

Lekawska-Andrinopoulou, L., Vasiliou, E.G., Georgakopoulos, D.G., Yialouris, C.P., and Georgiou, C.A. (2013). Rapid enzymatic method for pectin methyl esters determination. J Anal Methods Chem 2013, 854763.

Levesque-Tremblay, G., Muller, K., Mansfield, S.D., and Haughn, G.W. (2015a). HIGHLY METHYL ESTERIFIED SEEDS is a pectin methyl esterase involved in embryo development. Plant Physiol 167, 725–737.

Levesque-Tremblay, G., Pelloux, J., Braybrook, S.A., and Muller, K. (2015b). Tuning of pectin methylesterification, consequences for cell wall biomechanics and development. Planta 242, 791–811.

Li, T., Xu, Y., Zhang, L., Ji, Y., Tan, D., Yuan, H., and Wang, A. (2017). The Jasmonate-Activated Transcription Factor MdMYC2 Regulates ETHYLENE RESPONSE FACTOR and Ethylene Biosynthetic Genes to Promote Ethylene Biosynthesis during Apple Fruit Ripening. Plant Cell 29, 1316–1334.

Lionetti, V., Raiola, A., Camardella, L., Giovane, A., Obel, N., Pauly, M., Favaron, F., Cervone, F., and Bellincampi, D. (2007). Overexpression of pectin methylesterase inhibitors in Arabidopsis restricts fungal infection by Botrytis cinerea. Plant Physiol 143, 1871–1880.

Liu, W., Karemera, N.J.U., Wu, T., Yang, Y., Zhang, X., Xu, X., Wang, Y., and Han, Z. (2017). The ethylene response factor AtERF4 negatively regulates the iron deficiency response in Arabidopsis thaliana. PLoS One 12, e0186580.

Macquet, A., Ralet, M.C., Kronenberger, J., Marion-Poll, A., and North, H.M. (2007). In situ, chemical and macromolecular study of the composition of Arabidopsis thaliana seed coat mucilage. Plant Cell Physiol 48, 984–999.

Maruyama, Y., Yamoto, N., Suzuki, Y., Chiba, Y., Yamazaki, K., Sato, T., and Yamaguchi, J. (2013). The Arabidopsis transcriptional repressor ERF9 participates in resistance against necrotrophic fungi. Plant Sci 213, 79–87.

McGrath, K.C., Dombrecht, B., Manners, J.M., Schenk, P.M., Edgar, C.I., Maclean, D.J., Scheible, W.R., Udvardi, M.K., and Kazan, K. (2005). Repressor- and activator-type ethylene response factors functioning in jasmonate signaling and disease resistance identified via a genome-wide screen of Arabidopsis transcription factor gene expression. Plant Physiol 139, 949–959.

Micheli, F. (2001). Pectin methylesterases, cell wall enzymes with important roles in plant physiology. Trends Plant Sci 6, 414–419.

Muller, K., Levesque-Tremblay, G., Bartels, S., Weitbrecht, K., Wormit, A., Usadel, B., Haughn, G., and Kermode, A.R. (2013). Demethylesterification of cell wall pectins in Arabidopsis plays a role in seed germination. Plant Physiol 161, 305–316.

Nakano, T., Suzuki, K., Fujimura, T., and Shinshi, H. (2006). Genome-wide analysis of the ERF gene family in Arabidopsis and rice. Plant Physiol 140, 411–432.

Nguyen, H.P., Jeong, H.Y., Jeon, S.H., Kim, D., and Lee, H.P. (2017). Rice pectin methylesterase inhibitor28 (OsPMEI28) encodes afunctional PMEI and its overexpression results in a dwarf phenotypethrough increased pectin methylesterification levels. J Plant Physiol 208, 17–25.

Peaucelle, A., Braybrook, S.A., Le Guillou, L., Bron, E., Kuhlemeier, C., and Hofte, H. (2011). Pectin-induced changes in cell wall mechanics underlie organ initiation in Arabidopsis. Curr Biol 21, 1720–1726.

Pelletier, S., Orden, J.V., Wolf, S., Vissenberg, K., Delacourt, J., Ndong, Y.A., Pelloux, J., Bischoff, V., Urbrain, A., Mouille, G., Lemonnier, G., Renou, J.P., and Hofte, H. (2010). A role for pectin de-methylesterification in a developmentally regulated growth acceleration in darkgrown Arabidopsis hypocotyls. New Phytologist 188, 726–739.

Pelloux, J., Rusterucci, C., and Mellerowicz, E.J. (2007). New insights into pectin methylesterase structure and function. Trends Plant Sci 12, 267–277.

Ranocha, P., Francoz, E., Burlat, V., and Dunand, C. (2014). Expression of PRX36, PMEI6 and SBT1.7 is controlled by complex transcription factor regulatory networks for proper seed coat mucilage extrusion. Plant Signal Behav 9, e977734.

Rautengarten, C., Usadel, B., Neumetzler, L., Hartmann, J., Bussis, D., and Altmann, T. (2008). A subtilisin-like serine protease essential for mucilage release from Arabidopsis seed coats. Plant J 54, 466–480.

Reca, I.B., Lionetti, V., Camardella, L., D’Avino, R., Giardina, T., Cervone, F., and Bellincampi, D. (2012). A functional pectin methylesterase inhibitor protein (SolyPMEI) is expressed during tomato fruit ripening and interacts with PME-1. Plant Mol Biol 79, 429–442.

Riester, L., Koster-Hofmann, S., Doll, J., Berendzen, K.W., and Zentgraf, U. (2019). Impact of Alternatively Polyadenylated Isoforms of ETHYLENE RESPONSE FACTOR4 with Activator and Repressor Function on Senescence in Arabidopsis thaliana L. Genes (Basel) 10.

Rockel, N., Wolf, S., Kost, B., Rausch, T., and Greiner, S. (2008). Elaborate spatial patterning of cell-wall PME and PMEI at thepollen tube tip involves PMEI endocytosis, and reflects thedistribution of esterified and de-esterified pectins. The Plant J 53, 133–143.

Saez-Aguayo, S., Ralet, M.C., Berger, A., Botran, L., Ropartz, D., Marion-Poll, A., and North, H.M. (2013). PECTIN METHYLESTERASE INHIBITOR6 promotes Arabidopsis mucilage release by limiting methylesterification of homogalacturonan in seed coat epidermal cells. Plant Cell 25, 308–323.

Senechal, F., Mareck, A., Marcelo, P., Lerouge, P., and Pelloux, J. (2015). Arabidopsis PME17 Activity can be Controlled by Pectin Methylesterase Inhibitor4. Plant Signal Behav 10, e983351.

Shi, D., Ren, A., Tang, X., Qi, G., Xu, Z., Chai, G., Hu, R., Zhou, G., and Kong, Y. (2018). MYB52 Negatively Regulates Pectin Demethylesterification in Seed Coat Mucilage. Plant Physiol 176, 2737–2749.

Sullivan, S., Ralet, M.C., Berger, A., Diatloff, E., Bischoff, V., Gonneau, M., Marion-Poll, A., and North, H.M. (2011). CESA5 is required for the synthesis of cellulose with a role in structuring the adherent mucilage of Arabidopsis seeds. Plant Physiol 156, 1725–1739.

Turbant, A., Fournet, F., Lequart, M., Zabijak, L., Pageau, K., Bouton, S., and Van Wuytswinkel, O. (2016). PME58 plays a role in pectin distribution during seed coat mucilage extrusion through homogalacturonan modification. J Exp Bot 67, 2177–2190.

Voiniciuc, C., Dean, G.H., Griffiths, J.S., Kirchsteiger, K., Hwang, Y.T., Gillett, A., Dow, G., Western, T.L., Estelle, M., and Haughn, G.W. (2013). Flying saucer1 is a transmembrane RING E3 ubiquitin ligase that regulates the degree of pectin methylesterification in Arabidopsis seed mucilage. Plant Cell 25, 944–959.

Walker, M., Tehseen, M., Doblin, M.S., Pettolino, F.A., Wilson, S.M., Bacic, A., and Golz, J.F. (2011). The transcriptional regulator LEUNIG_HOMOLOG regulates mucilage release from the Arabidopsis testa. Plant Physiol 156, 46–60.

Wang, M., Xu, Z., Ahmed, R.I., Wang, Y., Hu, R., Zhou, G., and Kong, Y. (2019). Tubby-like Protein 2 regulates homogalacturonan biosynthesis in Arabidopsis seed coat mucilage. Plant Mol Biol 99, 421–436.

Wang, M., Yuan, D., Gao, W., Li, Y., Tan, J., and Zhang, X. (2013). A comparative genome analysis of PME and PMEI families reveals the evolution of pectin metabolism in plant cell walls. PLoS One 8, e72082.

Warde-Farley, D., Donaldson, S.L., Comes, O., Zuberi, K., Badrawi, R., Chao, P., Franz, M., Grouios, C., Kazi, F., Lopes, C.T., Mailand, A., Mostafavi, S., Montojo, J., Shao, Q., Wright, G., Bader, G.D., and Morris, Q. (2010). The GeneMANIA prediction server, biological network integration for gene prioritization and predicting gene function. Nucleic Acids Res 38, W214–220.

Western, T.L., Burn, J., Tan, W.L., Skinner, D.J., Martin-McCaffrey, L., Moffatt, B.A., and Haughn, G.W. (2001). Isolation and Characterization of Mutants Defective in Seed Coat Mucilage Secretory Cell Development in Arabidopsis. Plant Physiol 127, 998–1011.

Western, T.L., Skinner, D.J., and Haughn, G.W. (2000). Differentiation of mucilage secretory cells of the Arabidopsis seed coat. Plant Physiol 122, 345–355.

Willats, W.G.T., McCartney, L., and Knox, J.P. (2001a). In-situ analysis of pectic polysaccharides in seed mucilage and at the root surface of Arabidopsis thaliana. Planta 213, 37–44.

Willats, W.G.T., Orfila, C., Limberg, G., Buchholt, H.C., van Alebeek, G.-J.W.M., Voragen, A.G.J., Marcus, S.E., Christensen, T.M.I.E., Mikkelsen, J.D., Murray, B.S., and Knox, J.P. (2001b). Modulation of the Degree and Pattern of Methyl-esterification of Pectic Homogalacturonan in Plant Cell Walls. Journal of Biological Chemistry 276, 19404–19413.

Winter, D., Vinegar, B., Nahal, H., Ammar, R., Wilson, G.V., and Provart, N.J. (2007). An “Electronic Fluorescent Pictograph” browser for exploring and analyzing large-scale biological data sets. PLoS One 2, e718.

Wolf, S., Mouille, G., and Pelloux, J. (2009). Homogalacturonan methyl-esterification and plant development. Mol Plant 2, 851–860.

Wormit, A., and Usadel, B. (2018). The multifaced role of pectin methylesterase inhibitors. Int J Mol Sci 19, 2878.

Yang, Y., Nicolas, M., Zhang, J., Yu, H., Guo, D., Yuan, R., Zhang, T., Yang, J., Cubas, P., and Qin, G. (2018). The TIE1 transcriptional repressor controls shoot branching by directly repressing BRANCHED1 in Arabidopsis. PLoS Genet 14, e1007296.

Yang, Z., Tian, L., Latoszek-Green, M., Brown, D., and Wu, K. (2005). Arabidopsis ERF4 is a transcriptional repressor capable of modulating ethylene and abscisic acid responses. Plant Mol Biol 58, 585–596.

Yao, W., Wang, L., Wang, J., Ma, F., Yang, Y., Wang, C., Tong, W., Zhang, J., Xu, Y., Wang, X., Zhang, C., and Wang Y. (2017). VpPUB24, a novel gene from Chinese grapevine, Vitis pseudoreticulata, targets VpICE1 to enhance cold tolerance. J Exp Bot 68, 2933–2949.

Yuan, H., Meng, D., Gu, Z., Li, W., Wang, A., Yang, Q., Zhu, Y., and Li, T. (2014). A novel gene, MdSSK1, as a component of the SCF complex rather than MdSBP1 can mediate the ubiquitination of S-RNase in apple. J Exp Bot 65, 3121–3131.

Zhang, Z., Zhang, B., Chen, Z., Zhang, D., Zhang, H., Wang, H., Zhang, Y., Cai, D., Liu, J., Xiao, S., Huo, Y., Liu, J., Zhang, L., Wang, M., Liu, X., Xue, Y., Zhao, L., Zhou, Y., and Chen, H. (2018). A PECTIN METHYLESTERASE gene at the maize Ga1 locus confers male function in unilateral cross-incompatibility. Nat Commun 9, 3678.

Zhao, Y., Cheng, S., Song, Y., Huang, Y., Zhou, S., Liu, X., and Zhou, D.X. (2015). The Interaction between Rice ERF3 and WOX11 Promotes Crown Root Development by Regulating Gene Expression Involved in Cytokinin Signaling. Plant Cell 27, 2469–2483.

Zhou, X., Zhang, Z.L., Park, J., Tyler, L., Yusuke, J., Qiu, K., Nam, E.A., Lumba, S., Desveaux, D., McCourt, P., Kamiya, Y., and Sun, T.P. (2016). The ERF11 Transcription Factor Promotes Internode Elongation by Activating Gibberellin Biosynthesis and Signaling. Plant Physiol 171, 2760–2770.

